# Cell extrusion - a novel mechanism driving neural crest cell delamination

**DOI:** 10.1101/2024.03.09.584232

**Authors:** Emma Moore, Ruonan Zhao, Mary C McKinney, Kexi Yi, Christopher Wood, Paul Trainor

**Affiliations:** Stowers Institute for Medical Research, Kansas City, MO, USA; Department of Anatomy and Cell Biology, University of Kansas Medical Center, Kansas City, KS, USA

**Keywords:** neural crest cells, delamination, epithelial to mesenchymal transition, EMT, cell extrusion, Piezo1, mouse embryo

## Abstract

Neural crest cells (NCC) comprise a heterogeneous population of cells with variable potency, that contribute to nearly every tissue and organ system throughout the body. Considered unique to vertebrates, NCC are transiently generated within the dorsolateral region of the neural plate or neural tube, during neurulation. Their delamination and migration are crucial events in embryo development as the differentiation of NCC is heavily influenced by their final resting locations. Previous work in avian and aquatic species has shown that NCC delaminate via an epithelial-mesenchymal transition (EMT), which transforms these stem and progenitor cells from static polarized epithelial cells into migratory mesenchymal cells with fluid front and back polarity. However, the cellular and molecular drivers facilitating NCC delamination in mammals are poorly understood. We performed live timelapse imaging of NCC delamination in mouse embryos and discovered a group of cells that exit the neuroepithelium as isolated round cells, which then halt for a short period prior to acquiring the mesenchymal migratory morphology classically associated with most delaminating NCC. High magnification imaging and protein localization analyses of the cytoskeleton, together with measurements of pressure and tension of delaminating NCC and neighboring neuroepithelial cells, revealed these round NCC are extruded from the neuroepithelium prior to completion of EMT. Furthermore, we demonstrate that cranial NCC are extruded through activation of the mechanosensitive ion channel, PIEZO1, a key regulator of the live cell extrusion pathway, revealing a new role for PIEZO1 in neural crest cell development. Our results elucidating the cellular and molecular dynamics orchestrating NCC delamination support a model in which high pressure and tension in the neuroepithelium results in activation of the live cell extrusion pathway and delamination of a subpopulation of NCC in parallel with EMT. This model has broad implications for our understanding of cell delamination in development and disease.

## Introduction

Neural crest cells (NCC) are a multipotent migratory stem and progenitor cell population unique to vertebrates that are responsible for generating most of the craniofacial skeleton, peripheral nervous system and pigment cells (Le Douarin and Dupin 2018) (Bronner and LeDouarin 2012). NCC development can be divided into four distinct phases: formation, delamination, migration and differentiation (Mayor and Theveneau 2013). Delamination, the separation of tissue or individual cells from a surrounding tissue layer, occurs repeatedly throughout development, such as during gastrulation and neurulation, and in disease pathogenesis, such as in cancer invasion (Gilbert and Barresi 2016). Delamination is required for NCC to detach from the border region of the neural plate in which they form, and this occurs through an epithelial-mesenchymal transition (EMT) (Theveneau and Mayor 2012, Lee, Nagai et al. 2013).

EMT is a cell autonomous process in which an epithelial cell becomes mesenchymal through the loss of apical-basal polarity and cell-cell adhesions, and acquisition of front-back polarity, focal adhesions and migratory capacity (Acloque, Adams et al. 2009, Powell, Blasky et al. 2013, Yang, Antin et al. 2020, Zhao and Trainor 2023). Decades of research in avian and aquatic species have provided a rich understanding of the gene regulatory networks governing EMT during NCC delamination. Briefly, EMT master regulators *Zeb2*, *Snail1/2*, and *Twist1* are induced by WNT, NOTCH, TGFb and Hypoxia signaling pathways (Nieto, Sargent et al. 1994, LaBonne and Bronner-Fraser 2000, Carver, Jiang et al. 2001, Blanco, Barrallo-Gimeno et al. 2007, Acloque, Adams et al. 2009, Barriga, Maxwell et al. 2013, Powell, Blasky et al. 2013, Rogers, Saxena et al. 2013). In *Xenopus*, *snai1/2* are required and sufficient for NCC EMT and delamination. Similarly, both *Snai2* and *Zeb2* are required in chicken embryos (Nieto, Sargent et al. 1994, LaBonne and Bronner-Fraser 1998, Carl, Dufton et al. 1999, LaBonne and Bronner-Fraser 2000, del Barrio and Nieto 2002, Aybar, Nieto et al. 2003, Taneyhill, Coles et al. 2007). Expression of the EMT master regulators results in the repression of the epithelial cell-cell adhesion protein E-cadherin, and upregulation of the mesenchymal intermediate filament protein vimentin (Powell, Blasky et al. 2013).

*Zeb2*, *Snai1/2*, and *Twist1* are considered master regulators of EMT because of their universal roles in both development and disease associated EMT (Zheng and Kang 2014, Yang, Antin et al. 2020). However, *Zeb2*, *Snail1/2*, and *Twist1* loss-of-function among other genetic regulators of EMT, do not result in the perturbation of NCC delamination in mouse embryos (Chen and Behringer 1995, Van de Putte, Maruhashi et al. 2003, Murray and Gridley 2006, Van de Putte, Francis et al. 2007, Bildsoe, Loebel et al. 2009). This difference in mammalian regulation of NCC EMT may be attributable to variation in the spatiotemporal signaling. For example, unlike frog and chicken embryos, *Snai2* is not expressed in pre-migratory NCC in mouse (Sefton, Sanchez et al. 1998). In addition, *Twist1* expression in mouse NCC begins only after delamination has already occurred (Supplemental Figure 1) (Gitelman 1997). Another contributing factor may be functional redundancy between these key EMT regulators. However, redundancy has been difficult to discern in mouse since global knockout of *Snai1* results in embryonic lethality during gastrulation, which is prior to NCC formation and delamination (Carver, Jiang et al. 2001). Nonetheless, in conditional *Snai1/2* double mutant mice, although left-right asymmetry determination is disrupted, NCC formation and delamination are not affected. It is important to note however, that *Wnt1-Cre,* which is the gold standard in the field for NCC lineage and function studies, is activated too late to affect NCC EMT and delamination and this could explain the absence of an EMT associated phenotype following conditional deletion of EMT master regulators *Snai1*, Zeb2 and *Twist1* (Barriga, Trainor et al. 2015). Our basic understanding of the cellular dynamics of NCC delamination in mouse has also been hampered by the difficulty in visualizing developmental processes that occur during mouse embryo development *in utero* (Moore and Trainor 2022). Therefore, we still do not know what drives NCC delamination in mouse and thus further research is needed to improve our fundamental understanding of mammalian NCC development.

To address this gap in knowledge, we visualized the cellular dynamics of NCC delamination through time-lapse imaging of mouse embryo development. Consequently, we uncovered a previously undescribed population of delaminating NCC that were round and lacking in polarity and mesenchymal marker expression. These round cells differed considerably from the majority of delaminating NCC which exhibit a classic elongated morphology with front and back polarity. We therefore hypothesized these round NCC were delaminating by an alternative mechanism, specifically cell extrusion, in parallel with NCC that delaminate via EMT. Cell extrusion has been shown to reduce tissue stress caused by overcrowding by facilitating cell delamination. During cell extrusion, a cell is forcefully expelled from an epithelium by neighboring cells (Rosenblatt, Raff et al. 2001, Eisenhoffer, Loftus et al. 2012, Gudipaty, Lindblom et al. 2017, Ohsawa, Vaughen et al. 2018). Therefore, cell extrusion uniquely requires structural rearrangement of the cytoskeleton in both the delaminating and neighboring cells. Electron microscopy and immunostaining confirmed cell extrusion-specific cytoskeleton structures were present in round NCC and neighboring cells during delamination. Measurements of the relative internal pressure and edge tension in the neuroepithelium determined that regions undergoing NCC delamination exhibit higher tissue stress – as would be expected of cell extrusion. Activation of the cell extrusion pathway by mechanical induction is mediated through the mechanosensitive ion channel, PIEZO1 (Eisenhoffer, Loftus et al. 2012, Gudipaty, Lindblom et al. 2017), and single-cell RNA sequencing of the cranial region of mouse embryos exhibiting NCC delamination and migration together with immunostaining demonstrated PIEZO1 is expressed by NCC during their delamination. To elucidate whether NCC can delaminate be cell extrusion, embryos were cultured in the presence of an antagonist (GsMTx4) or agonist (Yoda1) of PIEZO1. Fewer delaminated NCC were observed following PIEZO1 inhibition, and furthermore these embryos lacked the round NCC population characteristic of cell extrusion. In contrast, when PIEZO1 was activated, the total number of delaminated NCC increased in association with the presence of round NCC.

Altogether, our data has revealed cell extrusion to be a novel mechanism by which NCC delaminate, and this cellular process occurs in parallel with EMT. These findings further our understanding of mammalian NCC development and have the potential to inform our knowledge of delamination in other biological contexts such as in cancer metastasis.

## Results

### A subpopulation of NCC delaminate as distinct round cells

To better define the cellular dynamics underpinning mammalian NCC delamination, we performed time-lapse imaging of cultured whole *Wnt1-Cre;R26R-mTmG* transgenic embryos. The *Wnt1-Cre;R26R-mTmG* transgenic line allows for lineage tracing of NCC beginning at their pre-migratory stage (Chai, Jiang et al. 2000). In these mice, *Cre*-mediated excision of a stop codon results in membrane tagged GFP expression and thus tissue-specific visualization of cellular morphology (Muzumdar, Tasic et al. 2007). Delamination occurs in a temporal wave beginning at around E8.5, or the 4-5 somite stage (Trainor and Tam, 1995), in the midbrain and expands anterior-posteriorly along the neural axis as the embryo develops. Therefore, to capture morphological changes from the beginning of delamination, imaging was focused on the region immediately posterior to the most recent domain of NCC delamination in the head at E8.5. The first cells to exit the neuroepithelium exhibit a round morphology that was maintained for 30 minutes (Figure 1A). This non-migratory round morphology was surprising as cells that undergo EMT acquire a front-back polarity and form filopodial protrusions that guide migration (Acloque, Adams et al. 2009, Powell, Blasky et al. 2013). Cells that delaminated following this 30-minute morphological transition were elongated and produced protrusions indicative of EMT completion (Figure 1A).

**Figure 1.**
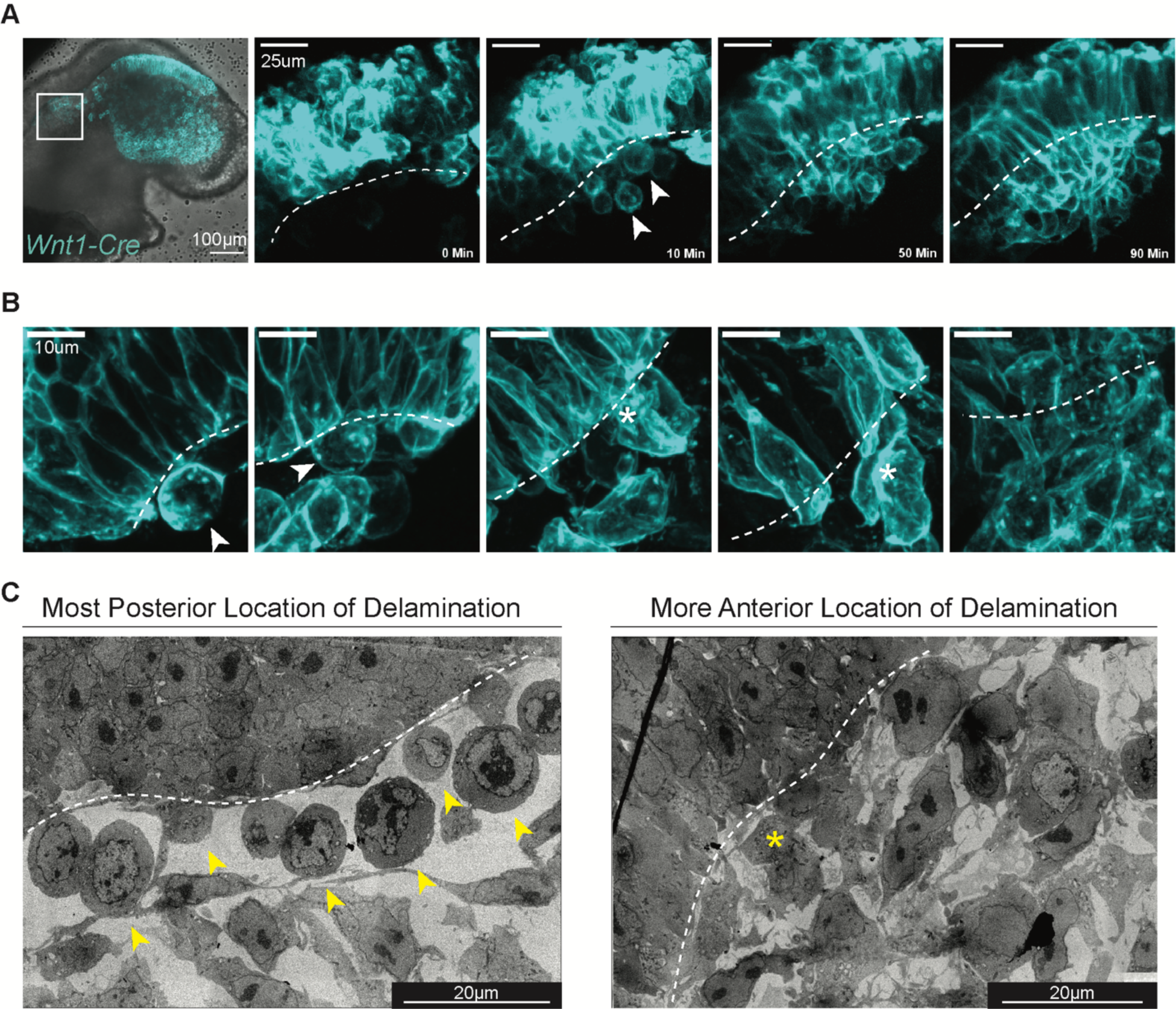
A subpopula1on of NCC delaminate as dis1nct round cells lacking polarity. **A**. Montage from -me-lapse imaging of a *Wnt1-Cre;R26R-mTmG* embryo at E8.5 shows the first NCC delaminate from the neuroepithelium as dis-nctly round cells. By 50 minutes the cells are elonga-ng and forming protrusions. At 90 minutes the elongated morphology is the only phenotype observed. **B**. Immunostaining of GFP (Cyan) on transverse histological sec-ons of *Wnt1-Cre;R26R-mTmG* embryos at E8.5 [not cultured] show the same cellular dynamics of delamina-ng NCC as compared to -me-lapse imaging. **C**. Transmission electron microscopy on sagiNal histological sec-ons of CD1 (wildtype) embryos at E8.5. More posterior loca-ons of the neuroepithelium exhibited a popula-on of round cells immediately outside of the basal edge whereas, more anterior regions, where delamina-on would have progressed for a longer -me, showed mesenchymal-like cells outside of the neuroepithelium. **A-C**. Arrow heads label the round subpopula-on, asterisks label presumably mesenchymal delamina-ng NCC and the dashed line outlines the basal edge of the neuroepithelium.

To determine if the round cells were indeed a novel delaminating population of NCC, and to rule out an artifact of culture and imaging conditions, we next sought to identify these round cells in non-cultured embryos *in situ*. Immunostaining for GFP on transverse histological sections of E8.5 *Wnt1-Cre;R26R-mTmG* embryos revealed a similar spectrum of cell morphologies (Figure 1B). Round NCC could be identified in select sections alongside elongated NCC, which comprised the predominant population of delaminated cells (Figure 1B). Evaluation of electron microscopy images from sagittal sections of E8.5 CD1 embryos also revealed the presence of round cells outside the neuroepithelium (Figure 1C). Notably, these round cells appeared to lack intracellular polarity. In more anterior (mature) locations, delaminating cells exhibited the classic mesenchymal morphology with distinct front-back polarity (Figure 1C). Altogether, our timelapse and static high magnification imaging of the dynamics of NCC delamination has uncovered a subpopulation of cells delaminating with a previously undescribed morphology, suggestive of a non-EMT mechanism, possibly cell extrusion.

### Cytoskeleton proteins are localized along round delaminating cells consistent with cell extrusion

We next wanted to understand why the dynamic morphology of these NCC differed from expected morphological changes typical of delamination by EMT. Breakdown of the basement membrane precedes NCC delamination and is necessary for emigration (Bronner-Fraser 1986, Duband and Thiery 1987, Bronner-Fraser and Lallier 1988, Coles, Gammill et al. 2006). To determine whether basement membrane breakdown had occurred when the round cells exit the neuroepithelium, we evaluated the expression of laminin – a major component of the basal lamina. Immunostaining of laminin indicated that the basement membrane had begun breaking down in concert with round cell delamination, as evidenced by lamina fragmentation near the cells (Supplemental Figure 2). There appeared to be more fragmented laminin in regions of elongated cells, consistent with subsequent EMT-mediated cell delamination and further progression of basement membrane breakdown (Supplemental Figure 2).

Since the basement membrane was broken, we then asked whether these round cells completed EMT and became mesenchymal, regardless of their initial atypical cellular morphology. Vimentin is an intermediate filament commonly used to identify mesenchymal cells following EMT (Usman, Waseem et al. 2021). Immunostaining for vimentin confirmed expression was present at the leading edge of elongated delaminating NCC, consistent with completion of their transition to a mesenchymal state (Figure 2A). In contrast, round NCC completely lacked vimentin activity, demonstrating that these cells were not yet mesenchymal (Figure 2A). These results indicated that the round NCC must be delaminating by a novel non-EMT mechanism.

**Figure 2.**
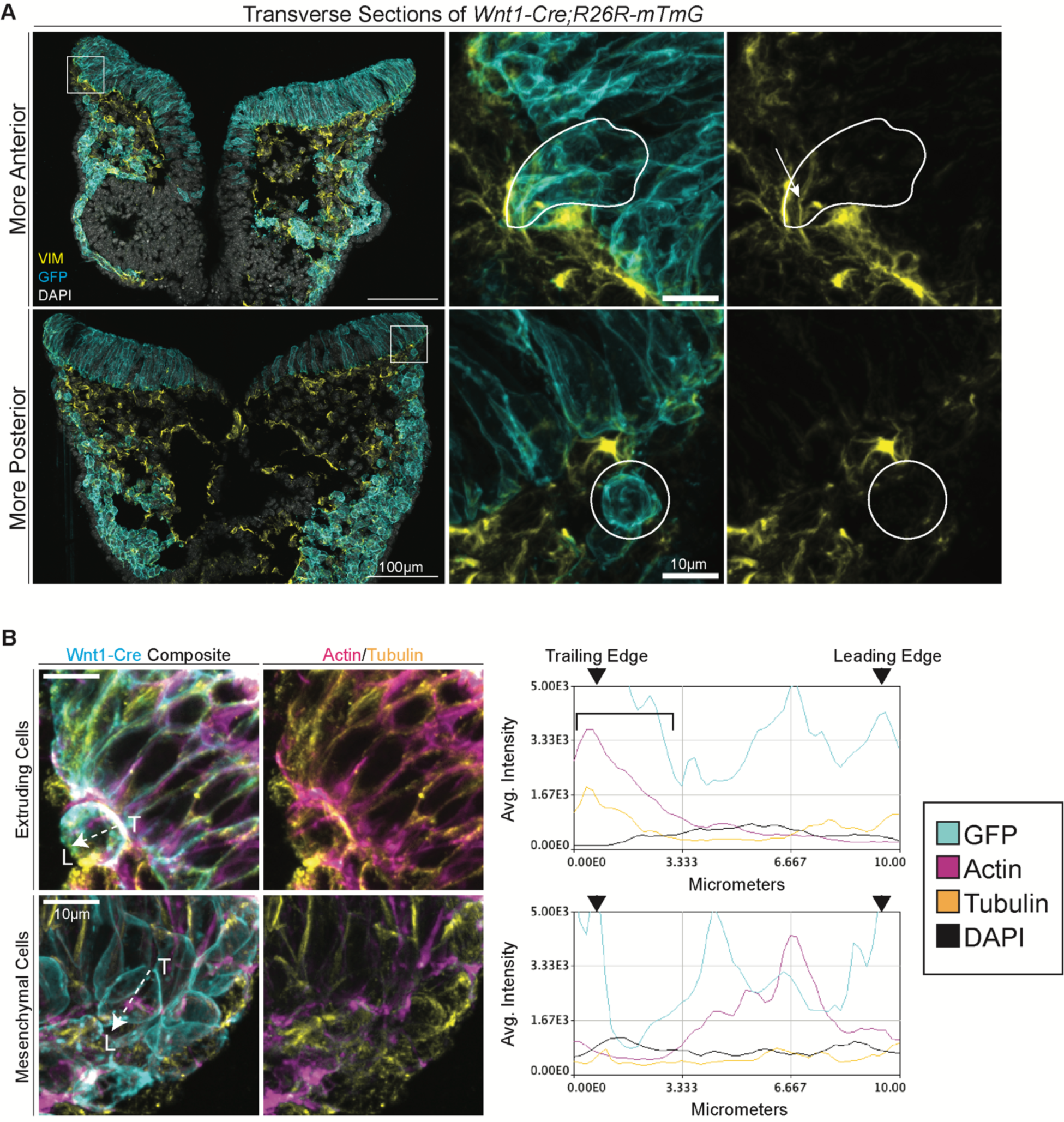
Cytoskeletal composition resembles cell extrusion. **A**. Immunostaining of Vimentin and GFP along with DAPI on transverse histological sections of *Wnt1-Cre;R26R-mTmG at E8.5*. The box indicates the location of the original tissue section from which the zoomed-in image was taken. The cells of interest are outlined in white with the top set of images showcasing NCC delaminating by EMT and the bottom set of images depicting an example of a round delaminating cell. The arrow is pointing out the expression of vimentin in the mesenchymal NCC which was not present in the round cell population. **B**. Immunostaining of Ac-n, Tubulin and GFP on transverse histological sec-ons of *Wnt1-Cre;R26R-mTmG* embryos at E8.5. Polyline kymograph analysis is presented in the graphs next to the cell from which the measurement was performed. The dashed line arrow indicates the region and direc-on the polyline was drawn across the cell of interest. T labels the trailing edge and L labels the leading edge. The bracket highlights the localiza-on of ac-n and tubulin found in the trailing edge of the extruding cell which was not present in the mesenchymal delamina-ng NCC.

Asymmetric cell division has previously been proposed as an alternative mechanism by which NCC could delaminate from the neuroepithelium (Erickson and Reedy 1998, Duband 2006, Ahlstrom and Erickson 2009). Furthermore, cells are known to acquire a round morphology when undergoing mitosis (Cadart, Zlotek-Zlotkiewicz et al. 2014). Cells undergoing division can be identified via electron microscopy by the condensation and alignment of chromosomes, and those that recently divided can be denoted by a reforming nuclear envelope filled with condensed chromatin (Supplemental Figure 3A). However, the round NCC delaminating from the neuroepithelium do not exhibit chromatin condensation (Figure 1C). Therefore, to conclusively discern whether the round cells were undergoing mitosis following delamination, we performed immunostaining with the G2/M marker phospho-histone H (pHH3). pHH3 was not observed in the round cell population indicating that the round morphology was not a result of, or associated with, cell division (Supplemental Figure 3B).

Cell extrusion is another form of delamination in which tissue stress triggers neighboring cells to expel adjacent cells from the epithelium in order to maintain tissue homeostasis (Eisenhoffer, Loftus et al. 2012, Marinari, Mehonic et al. 2012, Eisenhoffer and Rosenblatt 2013, Gudipaty, Lindblom et al. 2017, Gudipaty and Rosenblatt 2017, Ohsawa, Vaughen et al. 2018, Franco, Atieh et al. 2019). To facilitate delamination by cell extrusion, neighboring cells and the extruding cell undergo cytoskeletal reorganization. The cytoskeletal changes include formation of an actomyosin ring and orientation of microtubules along the cell membrane in the direction of expulsion (Rosenblatt, Raff et al. 2001, Slattum, McGee et al. 2009, Gu, Forostyan et al. 2011, Marshall, Lloyd et al. 2011, Slattum, Gu et al. 2014, Gudipaty, Lindblom et al. 2017, Gudipaty and Rosenblatt 2017, Ohsawa, Vaughen et al. 2018). The orchestration of inter and intracellular cytoskeleton changes during cell extrusion contrasts with EMT, which is considered a cell autonomous process. To determine whether these cytoskeletal differences could be identified, we analyzed the localization of actin and tubulin in the round and elongated NCC populations. Actin and tubulin were localized to the site of delamination, or trailing edge in round NCC as compared to other regions of the cell (Figure 2B). In NCC which delaminate by EMT, actin and tubulin were broadly distributed across the cell (Figure 2B). These observations were consistently observed across 12 round extruded NCC (localization in 8/12 cells, SEM=0.1361) and 13 EMT NCC (broad distribution in 11/13 cells, SEM=0.1001) compiled from 4 different embryos, leading to a p-value of 0.007742 by t-test analysis. Therefore, we hypothesize that some NCC delaminate by cell extrusion in parallel with EMT.

### The neuroepithelium contains higher tissue stress in regions of delamination

The cell extrusion pathway is activated by mechanical induction from physical tissue stress such as internal pressure and edge tension (Eisenhoffer, Loftus et al. 2012, Marinari, Mehonic et al. 2012, Eisenhoffer and Rosenblatt 2013, Franco, Atieh et al. 2019, Zulueta-Coarasa and Rosenblatt 2022). We reasoned that if cell extrusion could orchestrate NCC delamination, then higher pressure or tension should be evident in the dorsolateral most domain of the neuroepithelium from which NCC delaminate. The Cellular Force Inference Toolkit (CellFIT) enables the mapping of relative internal pressure and edge-edge tension within cells of a confluent tissue and has previously been used to study the role of tissue stress and cell extrusion in wound healing (Brodland, Veldhuis et al. 2014, Franco, Atieh et al. 2019). We used Tissue Analyzer to segment cells in transverse histological sections of 4-8 somite stage *Wnt1-Cre;R26R-mTmG* embryos (Figure 3A)(Aigouy, Umetsu et al. 2016). The segmented cells were then analyzed with CellFIT to calculate tissue stress. Higher internal pressure and edge-edge tension were consistently observed within the neuroepithelium in regions of NCC delamination (Figure 3B). Therefore, this suggests that mechanical tissue stress, which is required to induce cell extrusion, is present in the neuroepithelium during NCC delamination.

**Figure 3.**
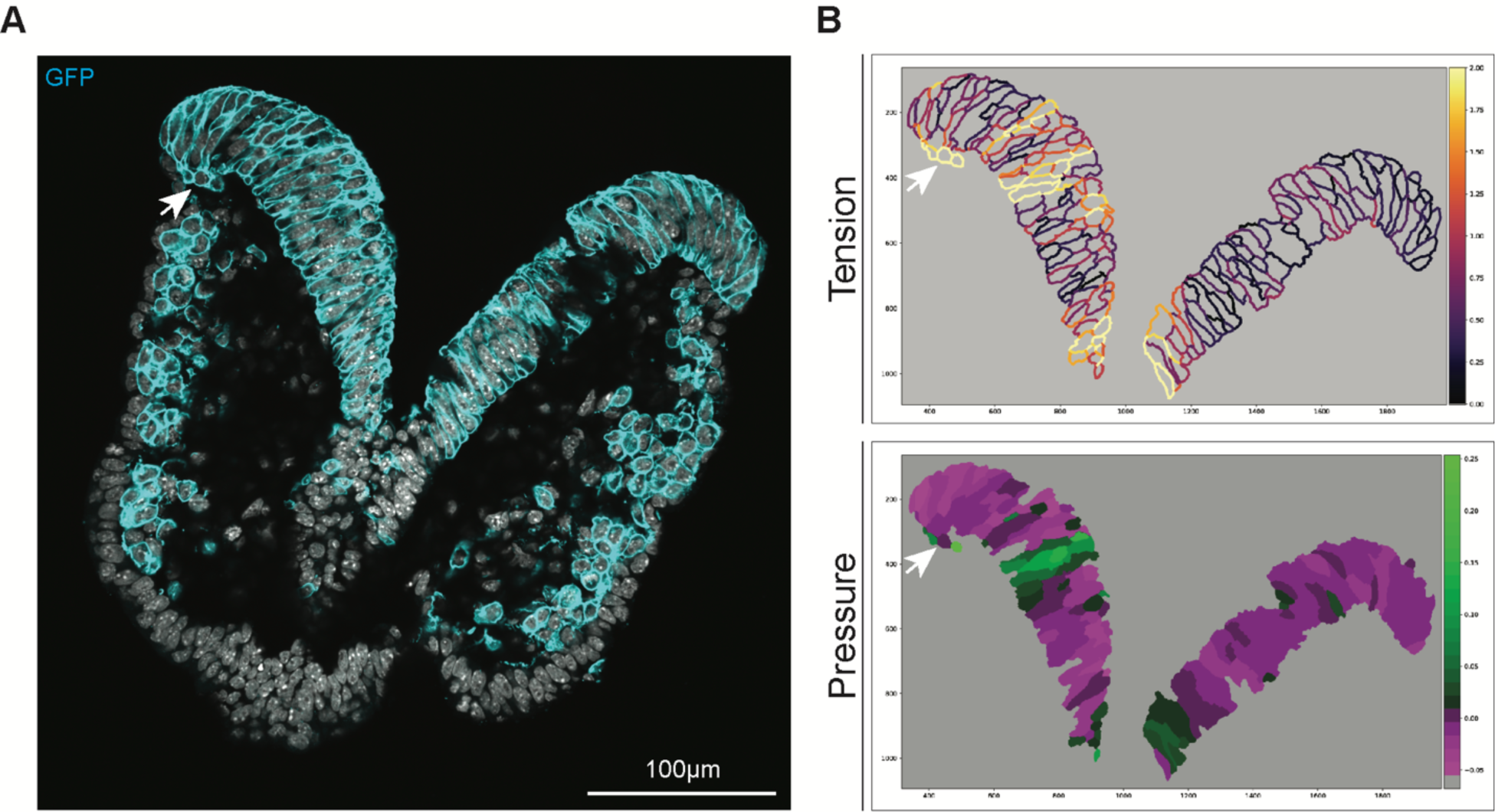
Delaminating NCC have higher edge-edge tension and internal pressure compared to surrounding neuroepithelial cells. **A**. Immunostaining of GFP along with DAPI staining on a transverse histological section of *Wnt1-Cre;R26R-mTmG* at E8.5, note this is only one z-slice. The arrow indicates 3 round cells delaminating from the neuroepithelium. **B**. Measurements of Tension and Pressure made by CellFIT analysis on segmented cells that originated from the tissue section in A. Higher edge tension (yellow) and internal pressure (green) are present in these delaminating NCC.

Cellular crowding has previously been linked to cases of cell extrusion as a contributing factor underpinning tissue stress (Eisenhoffer, Loftus et al. 2012, Marinari, Mehonic et al. 2012, Franco, Atieh et al. 2019). To determine whether the higher tissue stress associated with NCC delamination was due to overcrowding, we next measured the cell density across 3 distinct regions of the neuroepithelium, in 3 tissue sections at varying axial levels in 4-8 somite stage embryos. The 3 regions were defined as the dorsal most portion of the neuroepithelium from which NCC delaminate, the ventral region closest to the midline and the medial region that lies in between (Supplemental Figure 4). Cell density was measured in each region by quantification of the number of cells based on DAPI staining of nuclei using Cellpose (Stringer, Wang et al. 2021). Only the first 20-slices of the z-stack were used in each image to prevent overcounting. The total number of cells in the region was then divided by the volume of the 20-slice z-stack for normalization. No statistical difference was observed between the regions in the 9 sections analyzed (Supplemental Figure 4C). Delamination occurs in an anterior-posterior wave along the neuraxis. We reasoned that perhaps overcrowding was not present regionally but arises over developmental time. We therefore compared the cellular density from 3 sections each of 4, 6 and 8 somite stages embryos, however no statistical difference was observed in the cellular density between these 3 distinct developmental stages (Supplemental Figure 4D). These results suggest that overcrowding is not responsible for the tissue stress observed in the delaminating NCC. Therefore, the tissue stress must be caused by some other physical constraint, such as torsion of the neural folds during neurulation (Gilbert 2000, Nikolopoulou, Galea et al. 2017).

### Regulators of the cell extrusion pathway are expressed in pre-migratory NCC

Activation of the cell extrusion pathway by physical tissue stress is facilitated by the mechanosensitive ion channel, PIEZO1 (Eisenhoffer, Loftus et al. 2012, Gudipaty, Lindblom et al. 2017, Gudipaty and Rosenblatt 2017, Ohsawa, Vaughen et al. 2018). Although the complete downstream pathway following PIEZO1 activation remains to be elucidated (Gudipaty and Rosenblatt 2017), in apical extrusion it is known that activation of Piezo1 ultimately results in the production of sphingosine-1-phosphate (S1P) which then binds to its receptor S1P2 on neighboring cells (Gu, Forostyan et al. 2011, Eisenhoffer, Loftus et al. 2012, Gudipaty, Lindblom et al. 2017, Ohsawa, Vaughen et al. 2018). S1P2 activates Rho signaling through ARHGEF1 to target the localization of microtubules, actin and myosin within the neighboring cells (Slattum, McGee et al. 2009, Gu, Forostyan et al. 2011). To determine if these Piezo1 signaling components are expressed during NCC delamination, we analyzed our previously acquired single cell RNA-sequencing data of E8.5 mouse embryos which covered NCC development from pre-migratory to migratory stages (Falcon, Watt et al. 2022, Zhao, Moore et al. 2023). *Piezo1*, *S1p2* and *Arhgef1* are expressed throughout NCC development, including during delamination (Supplemental Figure 5), suggesting a potential role for Peizo1 mediated mechanotransduction in NCC delamination.

We next validated the expression of the extrusion pathway regulator, *Piezo1*, in pre-migratory cells within the neuroepithelium. We focused on *Piezo1* because its expression is specific to the extruding cell and live cell extrusion. Immunostaining indicated PIEZO1 is expressed in pre-migratory cells within the neuroepithelium prior to delamination (Figure 4). PIEZO1 was dynamically expressed in different regions of the same embryo, aligning with cell extrusion playing a temporally specific role in delamination (Figure 4A). Furthermore, PIEZO1 is known to localize in a cell at regions of cellular stress, and PIEZO1 was observed in the basal region of cells along the basal side of the neuroepithelium consistent with the detection of stress by CellFIT analyses (Figure 4B) (Ranade, Qiu et al. 2014, Gudipaty, Lindblom et al. 2017, Ellefsen, Holt et al. 2019, Holt, Zeng et al. 2021, Yao, Tijore et al. 2022). Overall, this data indicates components of the cell extrusion pathway are expressed in NCC during delamination.

**Figure 4.**
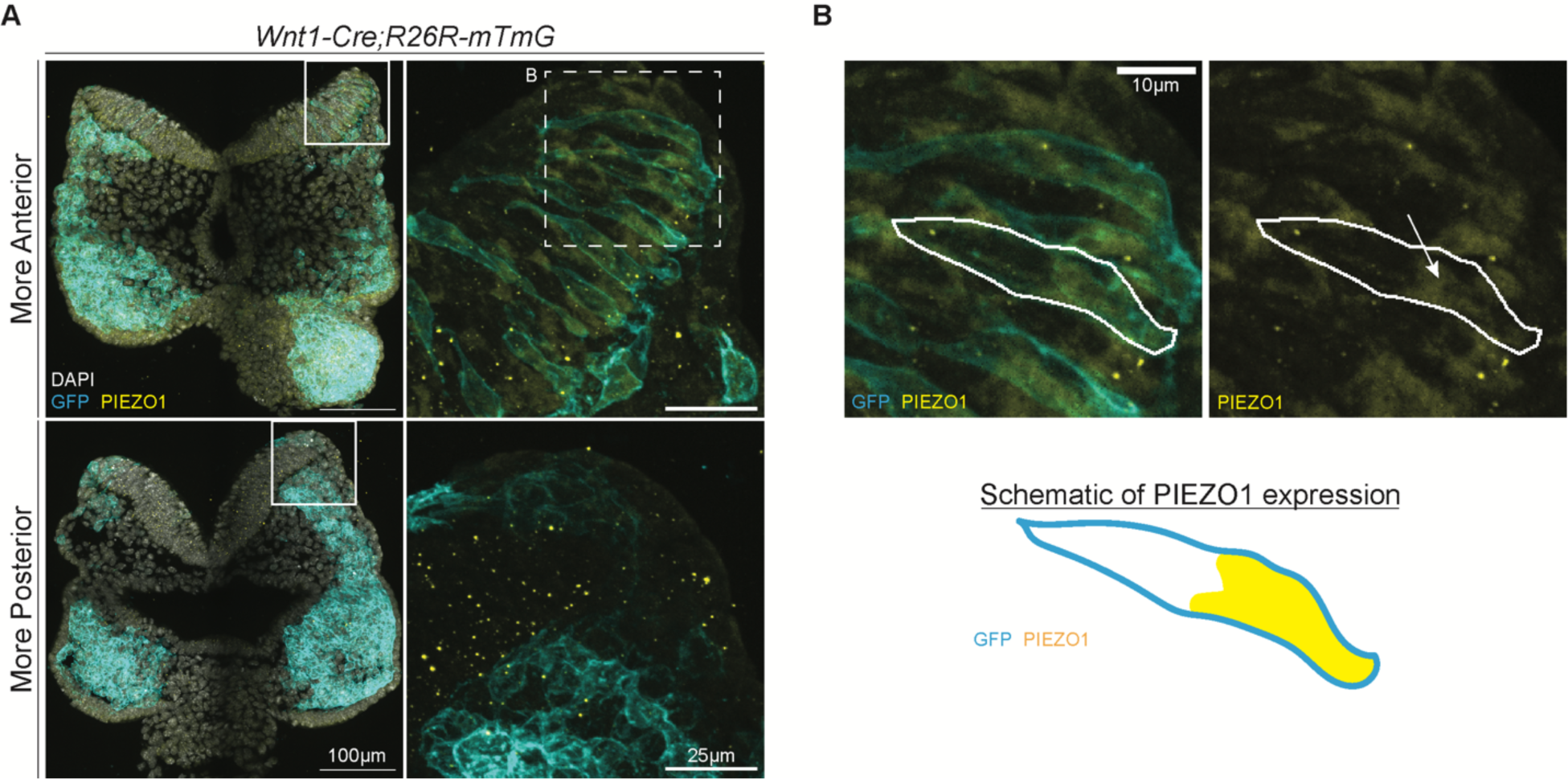
PIEZO1 is dynamically expressed in NCC during delamination. Immunostaining of PIEZO1 and GFP along with DAPI on transverse histological sections of *Wnt1-Cre;R26R-mTmG* taken from the same embryo at E8.5. **A**. The box indicates the location of the original tissue section from which the zoomed-in image on the right is taken. The expression of PIEZO1 can be found within GFP positive cells in the neuroepithelium as seen in the more anterior section. In the more posterior section where delamination is not occurring, PIEZO1 is not expressed in the neuroepithelium suggesting the expression of PIEZO1 to be highly dynamic. **B**. PIEZO1 expression is noticeably localized to the basal side of the cell within the neuroepithelium. These images are from the more anterior section of **A** as denoted by the dotted line box. A cell is outlined based on GFP expression to better display the basal localization. The arrow points towards the expression of PIEZO1 within the outlined cell. The schematic provides an additional representation of expression matching the outlined cell in the tissue section image.

### The cell extrusion pathway modulated delamination of the round NCC population

If PIEZO1 mediated cell extrusion is a mechanism by which NCC can delaminate, then modulating PIEZO1 activity should affect delamination and the number of migratory NCC. To test this model, CD1 embryos were cultured in the presence of a commonly used inhibitor of PIEZO1, GsMTx4, for 4 hours beginning at E8.25, which is prior to the start of cranial NCC delamination. Following embryo culture, the total number of SOX10 positive cells, a marker of migratory NCC, was quantified to determine how many NCC had delaminated (Figure 5A). The SOX10 population was then normalized to the total number of cells in the head according to DAPI staining. The GsMTx4 treated embryos displayed a significant decrease in NCC delamination as compared to vehicle-treated stage-matched littermates (Figure 5B). To confirm this decrease in delamination was due to perturbation of cell extrusion and thus loss of the round NCC population, this experiment was repeated with *Wnt1-Cre;R26R-mTmG* embryos. Treating E8.25 transgenic embryos with GsMTx4 completely diminished the presence of round NCC outside of the neuroepithelium at the most recent location of delamination as compared to vehicle-treated littermates (Figure 5C). This implies that PIEZO1 mediated mechanotransduction is required for NCC delamination via cell extrusion.

**Figure 5.**
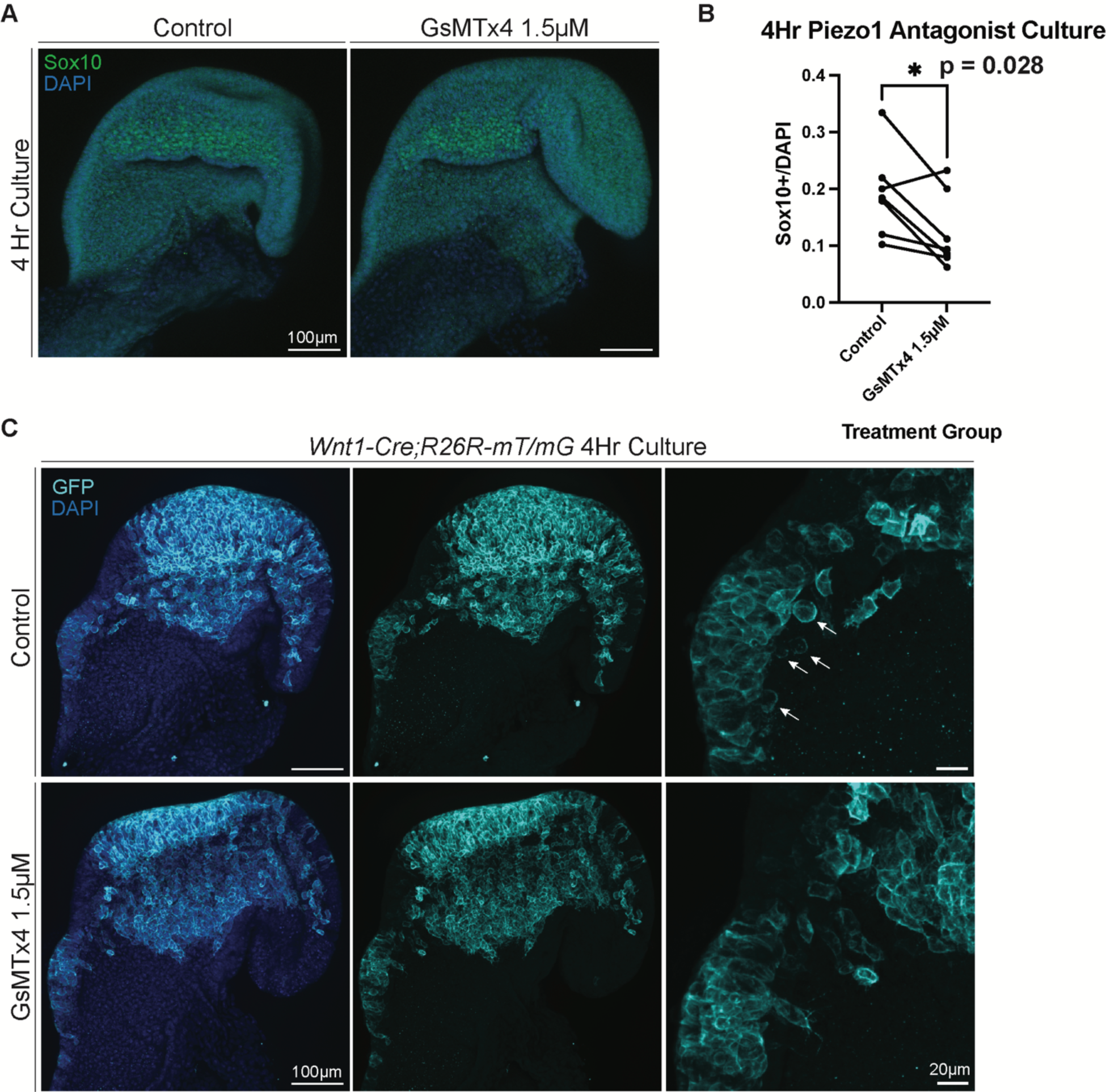
Inhibiting cell extrusion regulator, Piezo1, results in fewer migratory NCC and loss of extruded cell population. **A**. Immunostaining of SOX10 and DAPI on CD1 embryos at E8.5 following 4-hour culture with drug vehicle (control) or 1.5µM GsMTx4. **B**. Stage matched comparison between control and 1.5µM GsMTx4 embryos of the total number of migratory NCC (Sox10+) normalized to the total number of cells in the head (DAPI). A p value of 0.028 was determined by t-test suggesting NCC delamination decreases when PIEZO1 is inhibited by GsMTx4. **C**. Immunostaining of GFP and DAPI on *Wnt1-Cre;R26R-mTmG* embryos at E8.5 following 4-hour culture with drug vehicle (control) or 1.5µM GsMTx4. Arrows label the extruded cells present in the control embryos whereas no extruded cells were present in the PIEZO1 inhibitor treated embryos.

Consistent with this model, we then attempted to conversely enhance cell extrusion mediated NCC delamination via activation of PIEZO1 (Figure 5A) (Syeda, Xu et al. 2015). E8.25 CD1 embryos were treated with a specific PIEZO1 agonist, Yoda1, for 4 hours in culture, and we observed a significantly higher number of delaminated NCC as compared to vehicle-treated stage-matched littermates (Figure 6B). Furthermore, when E8.5 *Wnt1-Cre;R26R-mTmG* embryos were cultured in the presence of Yoda1, more NCC were observed outside the neuroepithelium at the most recent position of delamination (Figure 6C). Taken together, GsMTx4 antagonism and Yoda1 agonism of PIEZO1 activity indicates that some NCC can delaminate from the neuroepithelium by PIEZO1 induced cell extrusion, which can be identified and quantified by the presence of transiently round NCC.

**Figure 6.**
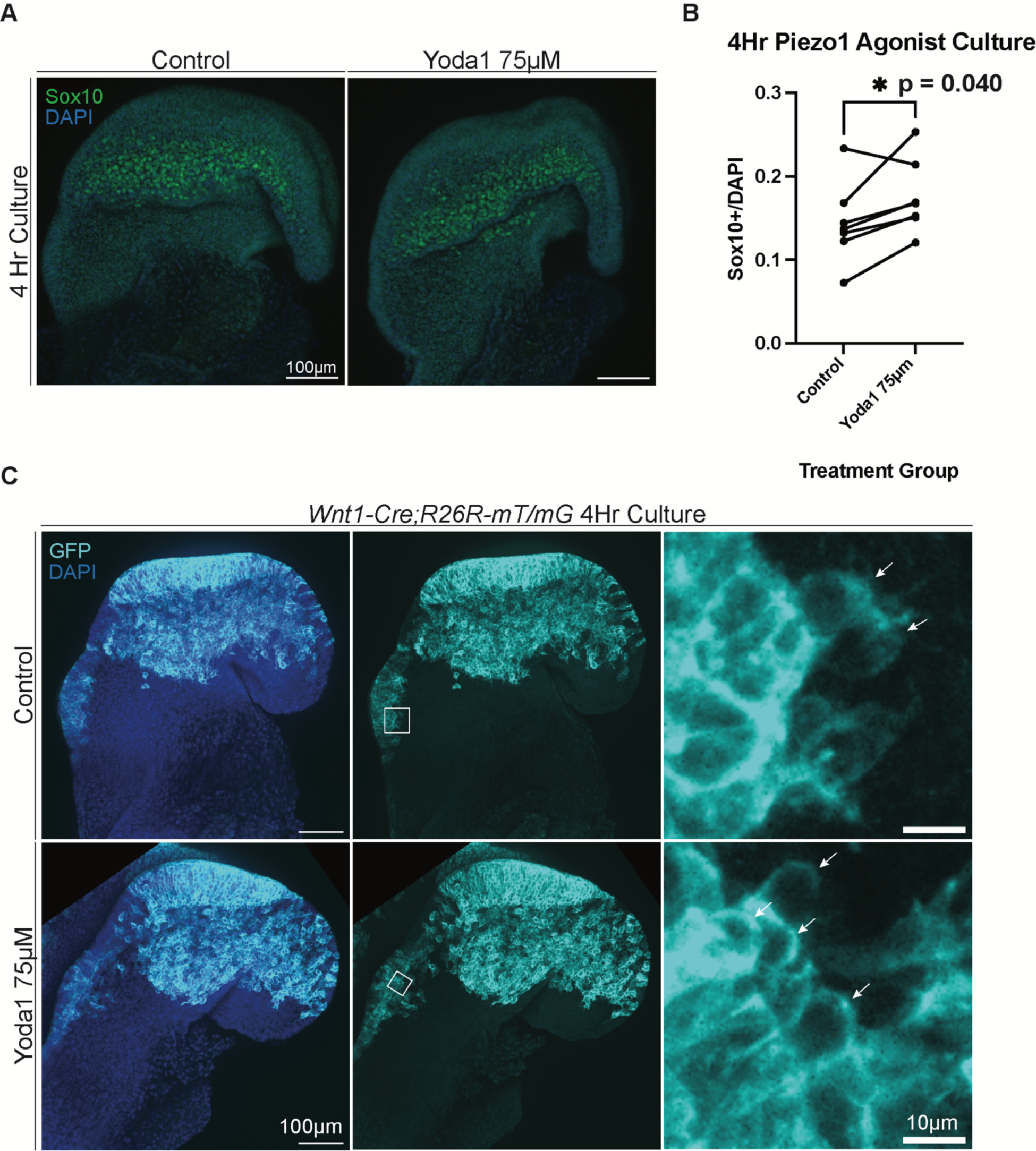
Activating cell extrusion regulator, Piezo1, increases the total number of migratory NCC. **A**. Immunostaining of Sox10 and DAPI on CD1 embryos at E8.5 following 4-hour culture with drug vehicle (control) or 75µM Yoda1. **B**. Stage matched comparison between control and 75µM Yoda1 embryos of the total number of migratory NCC (Sox10+) normalized to the total number of cells in the head (DAPI). A p value of 0.040 was determined by t-test suggesting NCC delamination increases when PIEZO1 is activated by Yoda1. **C**. Immunostaining of GFP and DAPI on *Wnt1-Cre;R26R-mTmG* embryos at E8.5 following 4-hour culture with drug vehicle (control) or 75µM Yoda1. Arrows label the extruded cells present in the control and PIEZO1 agonist treated embryos.

S1P and S1P2 signaling are considered downstream regulators of cell extrusion and we detected S1P kinase (SphK1) and S1P2 expression in NCC in our scRNA-seq data. However, it has previously been suggested that S1P signaling and S1P2 do not regulate basal cell extrusion events like NCC delamination. Cells that extrude basally have reduced levels of S1P compared to apical extruding cells (Gu, Forostyan et al. 2011). In addition, inhibiting S1P2 does not appear to prevent or induce basal extrusion (Gu, Forostyan et al. 2011, Yamamoto, Yako et al. 2016). Therefore, to test this idea and determine whether S1P signaling functions in NCC delamination, we cultured E8.25 CD1 embryos for 4 hours in the presence of the S1P2-specific inhibitor, JTE-013 (Supplemental Figure 6A). However, no measurable differences in the number of Sox10+ migrating NCC were found between the inhibitor-treated and vehicle-treated stage-matched littermates (Supplemental Figure 6B). Therefore, S1P2 does not facilitate NCC delamination, consistent with previous suggestions that S1P signaling does not regulate the process of basal cell extrusion.

## Discussion

Our results have elucidated a novel mechanism by which NCC delaminate during mammalian development. NCC undergo EMT and exit the neuroepithelium as mesenchymal cells, however, we have identified a subpopulation of NCC that delaminate by cell extrusion through PIEZO1 signaling, prior to transitioning to a mesenchymal state. Altogether, our data suggests the following cellular dynamics take place during NCC delamination in mammals: 1. Pressure and tension are high in cells along the basal edge or region of the neuroepithelium and the dorsolateral domain where NCC delaminate. 2. The ensuing tissue stress triggers PIEZO1 to induce the extrusion of a population of round NCC that have not yet completed EMT. 3. The remaining pre-migratory NCC transition to a mesenchymal state in concert with their delamination from the neuroepithelium (Figure 7).

**Figure 7.**
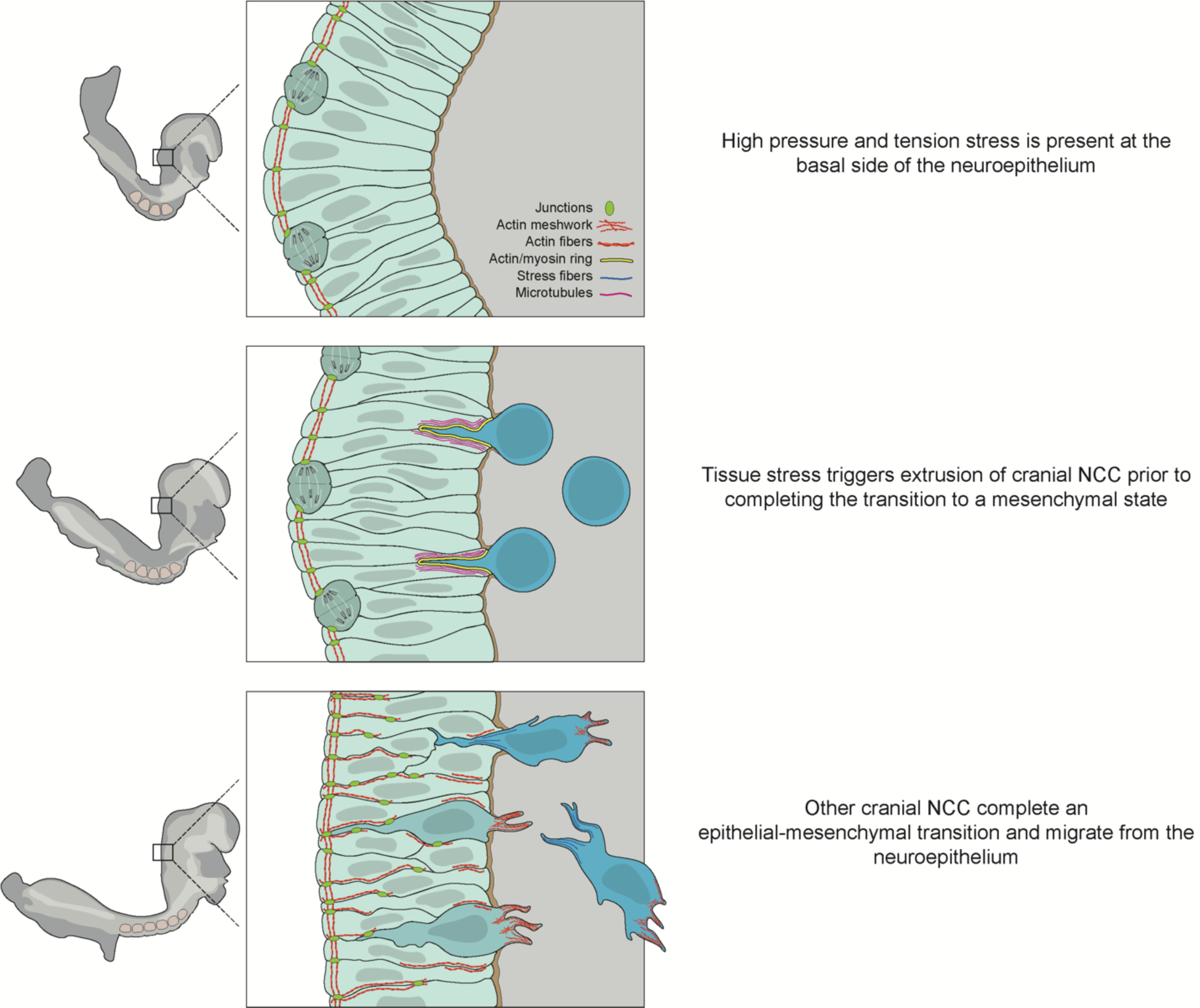
NCC delaminate by cell extrusion in parallel with EMT. An illustration of the proposed model for NCC delamination in mammalian development beginning with younger stages of development at the top and progressing to older stages while focused on the same region of the neuroepithelium. 1. Just prior to delamination there is a presence of high pressure and tension stress in the neuroepithelium. 2. Tissue stress triggers extrusion of NCC prior to the completion of EMT by activating PIEZO1. 3. Remaining NCC complete EMT and leave the neuroepithelium as mesenchymal migratory cells.

While our data indicates that NCC can delaminate by two different mechanisms, our previously published single cell RNA-seq data suggests the type of delamination does not restrict their capacity for differentiation. We previously identified two intermediate transition populations of NCC during their delamination, which were denoted as clusters 10 and 2 (Zhao, Moore et al. 2023). *Piezo1* is specifically expressed in cluster 2, indicating cluster 2 contains the extruding NCC population (Supplemental Figure 5E). Pseudo-time trajectory analysis performed on this single cell RNA-seq data determined that these intermediate transition populations are not lineage restricted (Zhao, Moore et al. 2023). Therefore, it is unlikely that the extruding cell population is limited to a certain fate at this developmental point. Furthermore, the delamination mechanism, cell extrusion versus EMT, does not appear to govern the ability of NCC to differentiate into their wide diversity of cell and tissues.

Time-lapse imaging and *in situ* identification of round cells at the onset of NCC delamination in a given axial region implicates cell extrusion as playing a potentially pivotal role in the initial exodus of NCC. These extruded NCC appear to be some of the first cells to break through the basement membrane and thus could act as leader cells that modify the surrounding extracellular matrix and communicate with follower cells. In the future, it will be interesting to define whether these extruded cells operate as leader cells as migration commences.

A significant question that arises from this work is: how many NCC delaminate by extrusion? Currently, the number of NCC that undergo cell extrusion compared to EMT cannot be determined because the round morphology is transient and there is no known marker to differentiate extrusion populations once this morphology has changed. More research is needed to identify a post-extrusion signature for long-term lineage tracing. Based on the number of NCC expressing *Piezo1* in the transition stage (cluster 2) and the reduction of delamination following inhibitor treatment, we estimate this number to be less than 30 percent (Supplemental Figure 5E). However, it is important to remember that this is still a significant difference when comparing the PIEZO1 agonist and antagonist treated embryos to controls.

Finally, the question remains as to whether cell extrusion is specific to mammalian development. We hypothesize NCC extrusion occurs in other model organisms based on the identification of round cells in other publications (Ahsan, Singh et al. 2019, Dunkel, Chaverra et al. 2020, Dobson, Barrell et al. 2023). For example, experiments in chicken embryos that focused on the role of FGFR1 in early trunk NCC migration noted non-polar round NCC that stall due to a loss of FGFR1 directed orientation (Dunkel, Chaverra et al. 2020). The timing between delamination and polarization of trunk NCC by FGFR1 signaling is consistent with the stalling of extruded cells observed in our time-lapse analyses. In zebrafish, Prickle1 loss-of-function results in persistent E-cadherin expression in migratory NCC, which is indicative of incomplete or hybrid EMT (Ahsan, Singh et al. 2019). Although these NCC expressed E-cadherin, delamination took place with a round morphology followed by blebbing (Ahsan, Singh et al. 2019). It is possible these cells delaminated by extrusion, which does not require complete downregulation of E-cadherin (Hogan, Dupre-Crochet et al. 2009, Lubkov and Bar-Sagi 2014, Wee, Hediyeh-Zadeh et al. 2020). While these studies in chicken and zebrafish align with our data on NCC extrusion, this cellular phenomenon needs to be evaluated more thoroughly to be conclusive and considered an evolutionarily conserved mechanism.

In summary, we have determined that cell extrusion plays an important role in mammalian cranial NCC delamination, implying it may also impact craniofacial development and the pathogenesis of craniofacial disorders. Consistent with this idea, previous research has shown cell extrusion facilitates secondary palatal shelf fusion (Kim, Lewis et al. 2015). Furthermore, two recently identified patients with biallelic mutations in *Piezo1*, present with distinct craniofacial anomalies including a flat facial gestalt and infraorbital hypoplasia (Lukacs, Mathur et al. 2015). These phenotypes are suggestive of a defect in neural crest cell development, perhaps a deficiency in cell extrusion leading to a reduction in the number of migratory NCC as was observed in our cultured mouse embryos in which PIEZO1 activity was inhibited. Continued studies will facilitate a better understanding of how *Piezo1* and cell extrusion impact later craniofacial development.

Cell extrusion is widely studied in the pathogenesis of cancer metastasis (Gudipaty and Rosenblatt 2017, Ohsawa, Vaughen et al. 2018). However, until now, the various extrusion models utilized in the field have focused on apical extrusion and the loss of key signaling components that result in a switch to basal extrusion (Slattum, McGee et al. 2009, Gu, Forostyan et al. 2011, Marshall, Lloyd et al. 2011, Slattum, Gu et al. 2014, Gudipaty and Rosenblatt 2017, Ohsawa, Vaughen et al. 2018). NCC delamination occurs via basal extrusion, providing an excellent model for further elucidation of basal-specific extrusion signaling mechanisms. Altogether, we have uncovered a novel mechanism driving mammalian NCC delamination, which furthers our understanding of NCC development with implications for craniofacial morphogenesis and cancer research.

## Materials and Methods

### Animal Husbandry

All animal experiments were performed in accordance with Stowers Institute for Medical Research Institutional Animal Care and Use Committee approved protocol #2022-014. Animals were housed in designated facilities on a 16-hour light and 8-hour dark cycle. Strains and transgenic lines used included CD1, FVB and *Wnt1-Cre;R26R-mT/mG*. *Wnt1-Cre* (H2afv^Tg(Wnt1-cre)11Rth^Tg(Wnt1-GAL4)11Rth/J) mice were obtained from The Jackson Laboratory (stock #003829) and maintained as male heterozygotes. Males were mated to female *R26R-mTmG* (Gt(ROSA)^26Sortm4(ACTB-tdTomato,-EGFP)Luo^/J) mice obtained from The Jackson Laboratory (stock #007576) to generate male *Wnt1-Cre;R26R-mT/mG* embryos.

### Embryo Collection

Timed mated females were checked for the presence of a vaginal plug and denoted embryonic day (E) 0.5. Embryos were collected on the designated day for the corresponding stage, following cervical dislocation of the mother. A detailed description of embryo dissection can be found in Chapter 18 “Live Imaging of the Dynamics of Mammalian Neural Crest Cell Migration” of Methods in Molecular Biology: Craniofacial Development (Moore and Trainor 2022).

### Time-lapse Imaging

Time-lapse imaging and image analysis were performed as previously described (Moore and Trainor 2022). In brief, embryos were cultured in 75% rat serum, 25% Fluorobrite DMEM with glutamax supplemented with 1% penicillin-streptomycin at 37 degrees Celsius in a 5% O2/5% CO2/N2 environment. Images were acquired on an inverted spinning disk confocal microscope (Nikon CSU-W1) with an sCMOS camera and piezo stage. Analysis was performed in Fiji and the 1µm z slices taken in each z-stack were max projected. To account for drift between z-slices and time points we ran StackReg and TurboReg respectively.

### Histology

Upon dissection, *Wnt1-Cre;R26R-mTmG* embryos were evaluated for the presence of GFP expressing cells indicating tissue specific Cre-mediated recombination has occurred in the dorsolateral neural plate which encompasses pre-migratory NCC. The embryos were then fixed in 4% PFA at 4 degrees Celsius overnight with rocking. The next day, the embryos were washed 3 times for 15 minutes each in 1X PBS with 0.1% Tween20 (PBSTw) and equilibrated in 30% sucrose in PBSTw at 4 degrees Celsius overnight with rocking. The next day the embryos were embedded in OCT and flash frozen on dry ice. Samples were cryosectioned at a thickness of 10-12 µm.

### Immunostaining

Transverse histological sections were washed 3 times for 5 minutes each with 1X PBS containing 0.3% Triton-X (PBSTx) to remove OCT. Blocking solution (varying serum concentrations in PBSTx) was applied to the sections for 2 hours at room temperature. Blocking solution was then replaced with fresh blocking solution containing the primary antibody and samples were left to incubate at 4°C overnight. Primary antibody solution was removed, and the samples were washed 3 times for 5 minutes each with PBSTx. Blocking solution with secondary antibody was then applied at a concentration of 1:500 along with 1:1000 DAPI and samples were left at room temperature for 2 hours in the dark. Finally, samples were washed 3 times for 5 minutes in PBSTx and mounted with a coverslip using Vectashield containing DAPI.

Vimentin (ab92547, Abcam) was used at a concentration of 1:500 with a blocking solution of 5% goat serum and 5% donkey serum. Laminin (ab11575, Abcam) was used at 1:200 with an 8% goat serum blocking solution. Alpha tubulin (T9026, Simga Aldrich) was used at 1:200 in combination with 1:500 phalloidin conjugated with 647 (Santa Cruz Biotechnology) in 8% goat serum and 2% Fab fragment donkey anti-mouse (715-007-003, Jackson ImmunoResearch) blocking solution. Piezo1 (NBP1-78446, Novus Biologicals) was used at 1:200 with 2% fetal bovine serum and 8% donkey serum. pHH3 (05-806, Sigma Millipore) was used at 1:500 with 3% BSA. All antibodies were applied together with 1:1000 GFP antibody (ab13970, Abcam) to visualize the GFP cell population.

Whole mount immunostaining of cultured embryos began with 4% PFA fixation at 4°C overnight with rocking. Embryos were then dehydrated into 100% methanol through a graded series of washes with increasing concentrations of methanol in PBSTw. Once in 100% methanol embryos were stored at −20°C overnight or until needed. The embryos were then bleached in a solution of 4:1:1 Methanol:DMSO:30% H2O2 respectively at room temperature for 2 hours. then rehydrated through a descending methanol in PBSTw series into PBSTw. The embryos were blocked in 10% fetal bovine serum in PBSTw for 2 hours at room temperature, and then incubated in fresh blocking solution containing 1:500 Sox10 antibody (ab155279, Abcam) at 4°C overnight with rocking. The next day the embryos were washed 5 times for 1 hour each before being incubated with secondary antibody at 1:500 together with 1:1000 DAPI in fresh blocking solution and rocked at 4°C overnight. The embryos were then washed 5 times for 1 hour and mounted in 0.8% low melting point agarose in PBSTw for imaging.

### Imaging of fixed tissue

Embryo sections were imaged at 40x, whereas *Wnt1-Cre;R26R-mTmG* whole mount embryos were imaged at 20x using a Nikon CSU-W1 inverted spinning disk confocal microscope equipped with a sCMOS camera and piezo stage. CD1 whole mount embryos were imaged at 10x with an LSM-800 Falcon inverted confocal microscope equipped with GaAsP detector and fast laser scanning technology.

### Electron Microscopy

CD1 embryos were collected at E8.5 around 6-8 somite stages and bisected along the sagittal plane. The CD1 embryos used to evaluate the general morphology of delaminating cells were dissected in 4% PFA before transfer to 2.5% glutaraldehyde, 2% PFA, 1mM CaCl3, 1g sucrose and 100mM sodium cacodylate at 4°C. Next, the samples were slowly dehydrated and embedded in resin. Sectioning was performed on an ultramicrotome and the tissue samples were imaged with a Technai Spirit Bio-TWIN 120 kV.

### Analysis of Cytoskeleton Protein Localization

To assess the localization of tubulin and actin in delaminating cells, sections of E8.5 *Wnt1-Cre;R26R-mTmG* embryos 4-8 somite stage embryos were immunostained with alpha-tubulin and phalloidin antibodies. Z-stack images of the 10-12µm sections (equivalent to one cell width) were max projected in FIJI. A 1-pixel width line was then drawn in the middle of the cell beginning at the edge of the cell towards the apical side of the neuroepithelium and ending at the basal side where the cell is no longer in the neuroepithelium. A polyline kymograph was plotted according to the line using “polyline kymograph jru v1” and plot colors were then matched to the false color of that assigned in the immunostaining image. The polyline kymograph plugin and further information can be found at: https://github.com/jayunruh/Jay_Plugins/blob/master/polyline_kymograph_jru_v1.java.

### Measuring Relative Tissue Stress

Transverse histological sections of *Wnt1-Cre;R26R-mTmG* embryos were immunostained for tubulin, phalloidin and GFP and imaged according to the methods outlined above. From the z-stack, a z-slice was chosen that displayed all the cells in the neuroepithelium. Sections were only 10µm or 1 cell width thick ensuring our z-slice was representative of cells in the same epithelial layer. The selected slice was placed in the FIJI plugin Tissue Analyzer (Aigouy, Umetsu et al. 2016). Images were segmented according to the recommendations of Tissue Analyzer authors and the settings resulting in the most consistent segmentation between samples – strong blur 10, weak blur 2, and removal of cells smaller than 10 pixels. False segmented regions and cells in areas outside the neuroepithelium were manually removed. The segmented images were then used to measure edge tension and internal pressure.

Measurements were performed by loading the segmentations in Zazu, otherwise known as 2D CellFIT (Brodland, Veldhuis et al. 2014). Meshes were created with 6 num nodes, or an average of 6 triple-junction nodes per cell edge, according to author recommendations and for consistency between samples (Brodland, Veldhuis et al. 2014). Tangent vectors were calculated using the nearest segment similar to what has previously been utilized in the field. Nearest segment provided the highest accuracy as it resulted in the lowest condition numbers and standard error, signs determined by the author to indicate an ideal setting. Output measurements of cells, edge tension and internal pressure were then mapped using Python. Measurements were performed for at least six different sections along the cranial region using four different samples from multiple litters.

### Measuring Cellular Density of the Neuroepithelium

Three embryos from different litters were analyzed for measurement of the cellular density of the neuroepithelium. Three sections were taken at varying axial levels within the cranial region of each embryo. Regions of interest (ROI) were drawn at the edge (dorsal), the center (medial) and ventral locations of each neural fold. The volume of each ROI was calculated in FIJI from the xyz resolution of the image. The first 20 slices of each z-stack were max-projected and then the total number of nuclei were counted based on DAPI staining using CellPose in FIJI (Stringer, Wang et al. 2021).

### Single Cell RNA-sequencing bioinformatic analyses

Bioinformatic analyses of single-cell RNA sequencing of E8.5 on *Mef2c-F10N-LacZ* and *Wnt1-Cre;R26R-eYFP* embryos were performed as previously described (Falcon, Watt et al. 2022, Zhao, Moore et al. 2023).

### Roller Culture and Drug Treatment

*Wnt1-Cre;R26R-mT/mG* and CD1 whole embryo roller culture was performed according to our previously published protocols (Sakai and Trainor 2014, Munoz and Trainor 2019, Moore and Trainor 2022). Embryos were cultured for 4 hours at 37°C in 50% rat serum/50% DMEM/F12 with glutamax and 1% penicillin-streptomycin in a 5% O2/5% CO2/N2 environment. Due to the variation in age that can occur within a litter, E8.0 embryos of the same litter were separated into 2 groups so that similar stages were represented in each and any unhealthy embryos or outliers in age were not utilized. One group received drug treatment (1.5µM GsMTx4 or 75µM Yoda1), while the other received only the vehicle in which the drug was dissolved as a control (ddH2O or DMSO respectively) (Bae, Sachs et al. 2011, Syeda, Xu et al. 2015, Gnanasambandam, Ghatak et al. 2017). Drug treatments were performed using three separate CD1 litters for Sox10 cell counting and two *Wnt1-Cre;R26R-mT/mG* litters with a total of three Cre+ embryos for each treatment condition and morphology assessment. Following culture, extraembryonic membranes were removed, and the embryos were imaged to confirm quality of health on a brightfield microscope. The embryos were then fixed in 4% PFA at 4°C overnight with rocking, and then processed for immunostaining or imaging as described above.

## Acknowledgements

The authors thank members of the Trainor laboratory for their constructive feedback and discussion on this project. We acknowledge our incredible animal technicians, Melissa Childers and Marina Thexton, as well as the Laboratory Animal Services facility at Stowers Institute for Medical Research for the animal care and husbandry that made this work possible. We are grateful to members of the Histology and Microscopy Cores for their assistance, training and insight. We thank Mark Miller who beautifully captured cell extrusion in his illustration in Figure 7. This research was made possible through funding provided by the National Institute for Dental and Craniofacial Research F31 DE032256 (E.L.M.), the Graduate School of the Stowers Institute for Medical Research (E.L.M.) and Stowers Institute for Medical Research (P.A.T).

## Notes

### Competing Interest Statement

The authors have declared no competing interest.

## References

Acloque, H., M. S. Adams, K. Fishwick, M. Bronner-Fraser and M. A. Nieto (2009). “Epithelial-mesenchymal transitions: the importance of changing cell state in development and disease.” J Clin Invest 119(6): 1438–1449.

Ahlstrom, J. D. and C. A. Erickson (2009). “The neural crest epithelial-mesenchymal transition in 4D: a ‘tail’ of multiple non-obligatory cellular mechanisms.” Development 136(11): 1801–1812.

Ahsan, K., N. Singh, M. Rocha, C. Huang and V. E. Prince (2019). “Prickle1 is required for EMT and migration of zebrafish cranial neural crest.” Dev Biol 448(1): 16–35.

Aigouy, B., D. Umetsu and S. Eaton (2016). “Segmentation and Quantitative Analysis of Epithelial Tissues.” Methods Mol Biol 1478: 227–239.

Aybar, M. J., M. A. Nieto and R. Mayor (2003). “Snail precedes slug in the genetic cascade required for the specification and migration of the Xenopus neural crest.” Development 130(3): 483–494.

Bae, C., F. Sachs and P. A. Gottlieb (2011). “The mechanosensitive ion channel Piezo1 is inhibited by the peptide GsMTx4.” Biochemistry 50(29): 6295–6300.

Barriga, E. H., P. H. Maxwell, A. E. Reyes and R. Mayor (2013). “The hypoxia factor Hif-1alpha controls neural crest chemotaxis and epithelial to mesenchymal transition.” J Cell Biol 201(5): 759–776.

Barriga, E. H., P. A. Trainor, M. Bronner and R. Mayor (2015). “Animal models for studying neural crest development: is the mouse different?” Development 142(9): 1555–1560.

Bildsoe, H., D. A. Loebel, V. J. Jones, Y. T. Chen, R. R. Behringer and P. P. Tam (2009). “Requirement for Twist1 in frontonasal and skull vault development in the mouse embryo.” Dev Biol 331(2): 176–188.

Blanco, M. J., A. Barrallo-Gimeno, H. Acloque, A. E. Reyes, M. Tada, M. L. Allende, R. Mayor and M. A. Nieto (2007). “Snail1a and Snail1b cooperate in the anterior migration of the axial mesendoderm in the zebrafish embryo.” Development 134(22): 4073–4081.

Brodland, G. W., J. H. Veldhuis, S. Kim, M. Perrone, D. Mashburn and M. S. Hutson (2014). “CellFIT: a cellular force-inference toolkit using curvilinear cell boundaries.” PLoS One 9(6): e99116.

Bronner, M. E. and N. M. LeDouarin (2012). “Development and evolution of the neural crest: an overview.” Dev Biol 366(1): 2–9.

Bronner-Fraser, M. (1986). “An antibody to a receptor for fibronectin and laminin perturbs cranial neural crest development in vivo.” Dev Biol 117(2): 528–536.

Bronner-Fraser, M. and T. Lallier (1988). “A monoclonal antibody against a laminin-heparan sulfate proteoglycan complex perturbs cranial neural crest migration in vivo.” J Cell Biol 106(4): 1321–1329.

Cadart, C., E. Zlotek-Zlotkiewicz, M. Le Berre, M. Piel and H. K. Matthews (2014). “Exploring the function of cell shape and size during mitosis.” Dev Cell 29(2): 159–169.

Carl, T. F., C. Dulon, J. Hanken and M. W. Klymkowsky (1999). “Inhibition of neural crest migration in Xenopus using antisense slug RNA.” Dev Biol 213(1): 101–115.

Carver, E. A., R. Jiang, Y. Lan, K. F. Oram and T. Gridley (2001). “The mouse snail gene encodes a key regulator of the epithelial-mesenchymal transition.” Mol Cell Biol 21(23): 8184–8188.

Chai, Y., X. Jiang, Y. Ito, P. Bringas, Jr., J. Han, D. H. Rowitch, P. Soriano, A. P. McMahon and H. M. Sucov (2000). “Fate of the mammalian cranial neural crest during tooth and mandibular morphogenesis.” Development 127(8): 1671–1679.

Chen, Z. F. and R. R. Behringer (1995). “twist is required in head mesenchyme for cranial neural tube morphogenesis.” Genes Dev 9(6): 686–699.

Coles, E. G., L. S. Gammill, J. H. Miner and M. Bronner-Fraser (2006). “Abnormalities in neural crest cell migration in laminin alpha5 mutant mice.” Dev Biol 289(1): 218–228.

del Barrio, M. G. and M. A. Nieto (2002). “Overexpression of Snail family members highlights their ability to promote chick neural crest formation.” Development 129(7): 1583–1593.

Dobson, L., W. B. Barrell, Z. Seraj, S. Lynham, S. Y. Wu, M. Krause and K. J. Liu (2023). “GSK3 and lamellipodin balance lamellipodial protrusions and focal adhesion maturation in mouse neural crest migration.” Cell Rep 42(9): 113030.

Duband, J. L. (2006). “Neural crest delamination and migration: integrating regulations of cell interactions, locomotion, survival and fate.” Adv Exp Med Biol 589: 45–77.

Duband, J. L. and J. P. Thiery (1987). “Distribution of laminin and collagens during avian neural crest development.” Development 101(3): 461–478.

Dunkel, H., M. Chaverra, R. Bradley and F. Lefcort (2020). “FGF signaling is required for chemokinesis and ventral migration of trunk neural crest cells.” Dev Dyn 249(9): 1077–1097.

Eisenhoffer, G. T., P. D. Lolus, M. Yoshigi, H. Otsuna, C. B. Chien, P. A. Morcos and J. Rosenblatt (2012). “Crowding induces live cell extrusion to maintain homeostatic cell numbers in epithelia.” Nature 484(7395): 546–549.

Eisenhoffer, G. T. and J. Rosenblatt (2013). “Bringing balance by force: live cell extrusion controls epithelial cell numbers.” Trends Cell Biol 23(4): 185–192.

Ellefsen, K. L., J. R. Holt, A. C. Chang, J. L. Nourse, J. Arulmoli, A. H. Mekhdjian, H. Abuwarda, F. Tombola, L. A. Flanagan, A. R. Dunn, I. Parker and M. M. Pathak (2019). “Myosin-II mediated traction forces evoke localized Piezo1-dependent Ca(2+) flickers.” Commun Biol 2: 298.

Erickson, C. A. and M. V. Reedy (1998). “Neural crest development: the interplay between morphogenesis and cell differentiation.” Curr Top Dev Biol 40: 177–209.

Falcon, K. T., K. E. N. Watt, S. Dash, R. Zhao, D. Sakai, E. L. Moore, S. Fitriasari, M. Childers, M. E. Sardiu, S. Swanson, D. Tsuchiya, J. Unruh, G. Bugarinovic, L. Li, R. Shiang, A. Achilleos, J. Dixon, M. J. Dixon and P. A. Trainor (2022). “Dynamic regulation and requirement for ribosomal RNA transcription during mammalian development.” Proc Natl Acad Sci U S A 119(31): e2116974119.

Franco, J. J., Y. Atieh, C. D. Bryan, K. M. Kwan and G. T. Eisenhoffer (2019). “Cellular crowding influences extrusion and proliferation to facilitate epithelial tissue repair.” Mol Biol Cell 30(16): 1890–1899.

Gilbert, S. F. (2000). Formation of the Neural Tube. Developmental Biology. Sunderland (MA), Sinauer Associates.

Gilbert, S. F. and M. J. F. Barresi (2016). Developmental biology. Sunderland, Massachusetts, U.S.A., Sinauer Associates, Inc., Publishers.

Gitelman, I. (1997). “Twist protein in mouse embryogenesis.” Dev Biol 189(2): 205–214.

Gnanasambandam, R., C. Ghatak, A. Yasmann, K. Nishizawa, F. Sachs, A. S. Ladokhin, S. I. Sukharev and T. M. Suchyna (2017). “GsMTx4: Mechanism of Inhibiting Mechanosensitive Ion Channels.” Biophys J 112(1): 31–45.

Gu, Y., T. Forostyan, R. Sabbadini and J. Rosenblatt (2011). “Epithelial cell extrusion requires the sphingosine-1-phosphate receptor 2 pathway.” J Cell Biol 193(4): 667–676.

Gudipaty, S. A., J. Lindblom, P. D. Lolus, M. J. Redd, K. Edes, C. F. Davey, V. Krishnegowda and J. Rosenblatt (2017). “Mechanical stretch triggers rapid epithelial cell division through Piezo1.” Nature 543(7643): 118–121.

Gudipaty, S. A. and J. Rosenblatt (2017). “Epithelial cell extrusion: Pathways and pathologies.” Semin Cell Dev Biol 67: 132–140.

Hogan, C., S. Dupre-Crochet, M. Norman, M. Kajita, C. Zimmermann, A. E. Pelling, E. Piddini, L. A. Baena-Lopez, J. P. Vincent, Y. Itoh, H. Hosoya, F. Pichaud and Y. Fujita (2009). “Characterization of the interface between normal and transformed epithelial cells.” Nat Cell Biol 11(4): 460–467.

Holt, J. R., W. Z. Zeng, E. L. Evans, S. H. Woo, S. Ma, H. Abuwarda, M. Loud, A. Patapoutian and M. M. Pathak (2021). “Spatiotemporal dynamics of PIEZO1 localization controls keratinocyte migration during wound healing.” Elife 10.

Kim, S., A. E. Lewis, V. Singh, X. Ma, R. Adelstein and J. O. Bush (2015). “Convergence and extrusion are required for normal fusion of the mammalian secondary palate.” PLoS Biol 13(4): e1002122.

LaBonne, C. and M. Bronner-Fraser (1998). “Neural crest induction in Xenopus: evidence for a two-signal model.” Development 125(13): 2403–2414.

LaBonne, C. and M. Bronner-Fraser (2000). “Snail-related transcriptional repressors are required in Xenopus for both the induction of the neural crest and its subsequent migration.” Dev Biol 221(1): 195–205.

Le Douarin, N. M. and E. Dupin (2018). “The “beginnings” of the neural crest.” Dev Biol 444 Suppl 1: S3–S13.

Lee, R. T., H. Nagai, Y. Nakaya, G. Sheng, P. A. Trainor, J. A. Weston and J. P. Thiery (2013). “Cell delamination in the mesencephalic neural fold and its implication for the origin of ectomesenchyme.” Development 140(24): 4890–4902.

Lubkov, V. and D. Bar-Sagi (2014). “E-cadherin-mediated cell coupling is required for apoptotic cell extrusion.” Curr Biol 24(8): 868–874.

Lukacs, V., J. Mathur, R. Mao, P. Bayrak-Toydemir, M. Procter, S. M. Cahalan, H. J. Kim, M. Bandell, N. Longo, R. W. Day, D. A. Stevenson, A. Patapoutian and B. L. Krock (2015). “Impaired PIEZO1 function in patients with a novel autosomal recessive congenital lymphatic dysplasia.” Nat Commun 6: 8329.

Marinari, E., A. Mehonic, S. Curran, J. Gale, T. Duke and B. Baum (2012). “Live-cell delamination counterbalances epithelial growth to limit tissue overcrowding.” Nature 484(7395): 542–545.

Marshall, T. W., I. E. Lloyd, J. M. Delalande, I. Nathke and J. Rosenblatt (2011). “The tumor suppressor adenomatous polyposis coli controls the direction in which a cell extrudes from an epithelium.” Mol Biol Cell 22(21): 3962–3970.

Mayor, R. and E. Theveneau (2013). “The neural crest.” Development 140(11): 2247–2251.

Moore, E. L. and P. A. Trainor (2022). “Live Imaging of the Dynamics of Mammalian Neural Crest Cell Migration.” Methods Mol Biol 2403: 263–276.

Munoz, W. A. and P. A. Trainor (2019). “Mouse Embryo Culture for the Study of Neural Crest Cells.” Methods Mol Biol 1976: 107–119.

Murray, S. A. and T. Gridley (2006). “Snail family genes are required for lel-right asymmetry determination, but not neural crest formation, in mice.” Proc Natl Acad Sci U S A 103(27): 10300–10304.

Muzumdar, M. D., B. Tasic, K. Miyamichi, L. Li and L. Luo (2007). “A global double-fluorescent Cre reporter mouse.” Genesis 45(9): 593–605.

Nieto, M. A., M. G. Sargent, D. G. Wilkinson and J. Cooke (1994). “Control of cell behavior during vertebrate development by Slug, a zinc finger gene.” Science 264(5160): 835–839.

Nikolopoulou, E., G. L. Galea, A. Rolo, N. D. Greene and A. J. Copp (2017). “Neural tube closure: cellular, molecular and biomechanical mechanisms.” Development 144(4): 552–566.

Ohsawa, S., J. Vaughen and T. Igaki (2018). “Cell Extrusion: A Stress-Responsive Force for Good or Evil in Epithelial Homeostasis.” Dev Cell 44(4): 532.

Powell, D. R., A. J. Blasky, S. G. Britt and K. B. Artinger (2013). “Riding the crest of the wave: parallels between the neural crest and cancer in epithelial-to-mesenchymal transition and migration.” Wiley Interdiscip Rev Syst Biol Med 5(4): 511–522.

Ranade, S. S., Z. Qiu, S. H. Woo, S. S. Hur, S. E. Murthy, S. M. Cahalan, J. Xu, J. Mathur, M. Bandell, B. Coste, Y. S. Li, S. Chien and A. Patapoutian (2014). “Piezo1, a mechanically activated ion channel, is required for vascular development in mice.” Proc Natl Acad Sci U S A 111(28): 10347–10352.

Rogers, C. D., A. Saxena and M. E. Bronner (2013). “Sip1 mediates an E-cadherin-to-N-cadherin switch during cranial neural crest EMT.” J Cell Biol 203(5): 835–847.

Rosenblatt, J., M. C. Raff and L. P. Cramer (2001). “An epithelial cell destined for apoptosis signals its neighbors to extrude it by an actin-and myosin-dependent mechanism.” Curr Biol 11(23): 1847–1857.

Sakai, D. and P. A. Trainor (2014). “Gene transfer techniques in whole embryo cultured post-implantation mouse embryos.” Methods Mol Biol 1092: 227–234.

Selon, M., S. Sanchez and M. A. Nieto (1998). “Conserved and divergent roles for members of the Snail family of transcription factors in the chick and mouse embryo.” Development 125(16): 3111–3121.

Slattum, G., Y. Gu, R. Sabbadini and J. Rosenblatt (2014). “Autophagy in oncogenic K-Ras promotes basal extrusion of epithelial cells by degrading S1P.” Curr Biol 24(1): 19–28.

Slattum, G., K. M. McGee and J. Rosenblatt (2009). “P115 RhoGEF and microtubules decide the direction apoptotic cells extrude from an epithelium.” J Cell Biol 186(5): 693–702.

Stringer, C., T. Wang, M. Michaelos and M. Pachitariu (2021). “Cellpose: a generalist algorithm for cellular segmentation.” Nat Methods 18(1): 100–106.

Syeda, R., J. Xu, A. E. Dubin, B. Coste, J. Mathur, T. Huynh, J. Matzen, J. Lao, D. C. Tully, I. H. Engels, H. M. Petrassi, A. M. Schumacher, M. Montal, M. Bandell and A. Patapoutian (2015). “Chemical activation of the mechanotransduction channel Piezo1.” Elife 4.

Taneyhill, L. A., E. G. Coles and M. Bronner-Fraser (2007). “Snail2 directly represses cadherin6B during epithelial-to-mesenchymal transitions of the neural crest.” Development 134(8): 1481–1490.

Theveneau, E. and R. Mayor (2012). “Neural crest delamination and migration: from epithelium-to-mesenchyme transition to collective cell migration.” Dev Biol 366(1): 34–54.

Usman, S., N. H. Waseem, T. K. N. Nguyen, S. Mohsin, A. Jamal, M. T. Teh and A. Waseem (2021). “Vimentin Is at the Heart of Epithelial Mesenchymal Transition (EMT) Mediated Metastasis.” Cancers (Basel) 13(19).

Van de Putte, T., A. Francis, L. Nelles, L. A. van Grunsven and D. Huylebroeck (2007). “Neural crest-specific removal of Zpx1b in mouse leads to a wide range of neurocristopathies reminiscent of Mowat-Wilson syndrome.” Hum Mol Genet 16(12): 1423–1436.

Van de Putte, T., M. Maruhashi, A. Francis, L. Nelles, H. Kondoh, D. Huylebroeck and Y. Higashi (2003). “Mice lacking ZFHX1B, the gene that codes for Smad-interacting protein-1, reveal a role for multiple neural crest cell defects in the etiology of Hirschsprung disease-mental retardation syndrome.” Am J Hum Genet 72(2): 465–470.

Wee, K., S. Hediyeh-Zadeh, K. Duszyc, S. Verma, N. N. B. S. Khare, A. Varma, R. J. Daly, A. S. Yap, M. J. Davis and S. Budnar (2020). “Snail induces epithelial cell extrusion by regulating RhoA contractile signalling and cell-matrix adhesion.” J Cell Sci 133(13).

Yamamoto, S., Y. Yako, Y. Fujioka, M. Kajita, T. Kameyama, S. Kon, S. Ishikawa, Y. Ohba, Y. Ohno, A. Kihara and Y. Fujita (2016). “A role of the sphingosine-1-phosphate (S1P)-S1P receptor 2 pathway in epithelial defense against cancer (EDAC).” Mol Biol Cell 27(3): 491–499.

Yang, J., P. Antin, G. Berx, C. Blanpain, T. Brabletz, M. Bronner, K. Campbell, A. Cano, J. Casanova, G. Christofori, S. Dedhar, R. Derynck, H. L. Ford, J. Fuxe, A. Garcia de Herreros, G. J. Goodall, A. K. Hadjantonakis, R. Y. J. Huang, C. Kalcheim, R. Kalluri, Y. Kang, Y. Khew-Goodall, H. Levine, J. Liu, G. D. Longmore, S. A. Mani, J. Massague, R. Mayor, D. McClay, K. E. Mostov, D. F. Newgreen, M. A. Nieto, A. Puisieux, R. Runyan, P. Savagner, B. Stanger, M. P. Stemmler, Y. Takahashi, M. Takeichi, E. Theveneau, J. P. Thiery, E. W. Thompson, R. A. Weinberg, E. D. Williams, J. Xing, B. P. Zhou, G. Sheng and E. M. T. I. Association (2020). “Guidelines and definitions for research on epithelial-mesenchymal transition.” Nat Rev Mol Cell Biol 21(6): 341–352.

Yao, M., A. Tijore, D. Cheng, J. V. Li, A. Hariharan, B. Martinac, G. Tran Van Nhieu, C. D. Cox and M. Sheetz (2022). “Force-and cell state-dependent recruitment of Piezo1 drives focal adhesion dynamics and calcium entry.” Sci Adv 8(45): eabo1461.

Zhao, R., E. L. Moore, M. M. Gogol, J. R. Uhruh, Z. Yu, A. Scott, Y. Wang, N. K. Rajendran and P. A. Trainor (2023). “Identification and characterization of intermediate states in mammalian neural crest cell epithelial to mesenchymal transition and delamination.” bioRxiv.

Zhao, R. and P. A. Trainor (2023). “Epithelial to mesenchymal transition during mammalian neural crest cell delamination.” Semin Cell Dev Biol 138: 54–67.

Zheng, H. and Y. Kang (2014). “Multilayer control of the EMT master regulators.” Oncogene 33(14): 1755–1763.

Zulueta-Coarasa, T. and J. Rosenblatt (2022). “The role of tissue maturity and mechanical state in controlling cell extrusion.” Curr Opin Genet Dev 72: 1–7.

