## Supplemental Figures for "Cell extrusion - a novel mechanism driving neural crest cell delamination"

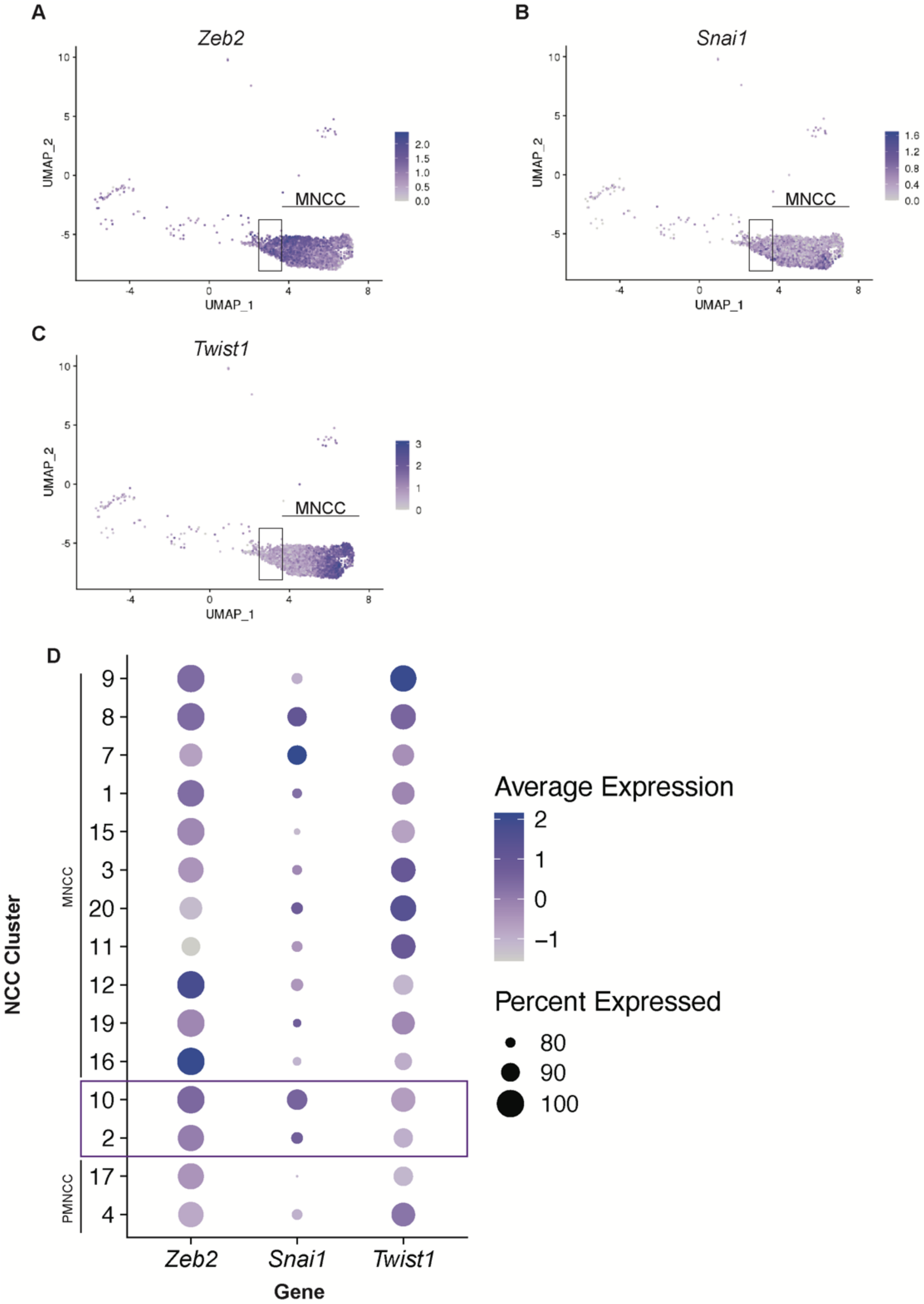


**Supplemental Figure 1. Master regulators of EMT are expressed at different stages of NCC development in mouse.** Single cell RNA-sequencing of the cranial region of *Wnt1-Cre;R26R-eYFP* and *Mef2c-F10n-LacZ* embryos at E8.5. The box indicates the delaminating NCC population. Migratory NCC are denoted by the label MNCC. PMNCC labels the pre-migratory NCC populations. **A-C**. UMAP expression of *Zeb2*, *Snai1* and *Twist1* in NCC across all stages of development from preimplantation to migration and early differentiation. **A**. *Zeb2* is highly expressed in the transition and early migratory NCC. **B**. *Snai1* is broadly expressed along the entire NCC cluster. **C.** *Twist1* is enriched in the late migratory NCC population. **D**. Dot plot of *Zeb2*, *Snai1* and *Twist1* expression at each state of NCC development. *Zeb2* and *Snai1* have an increased expression in the transition population as compared to PMNCC. *Twist1* is enriched in the MNCC.

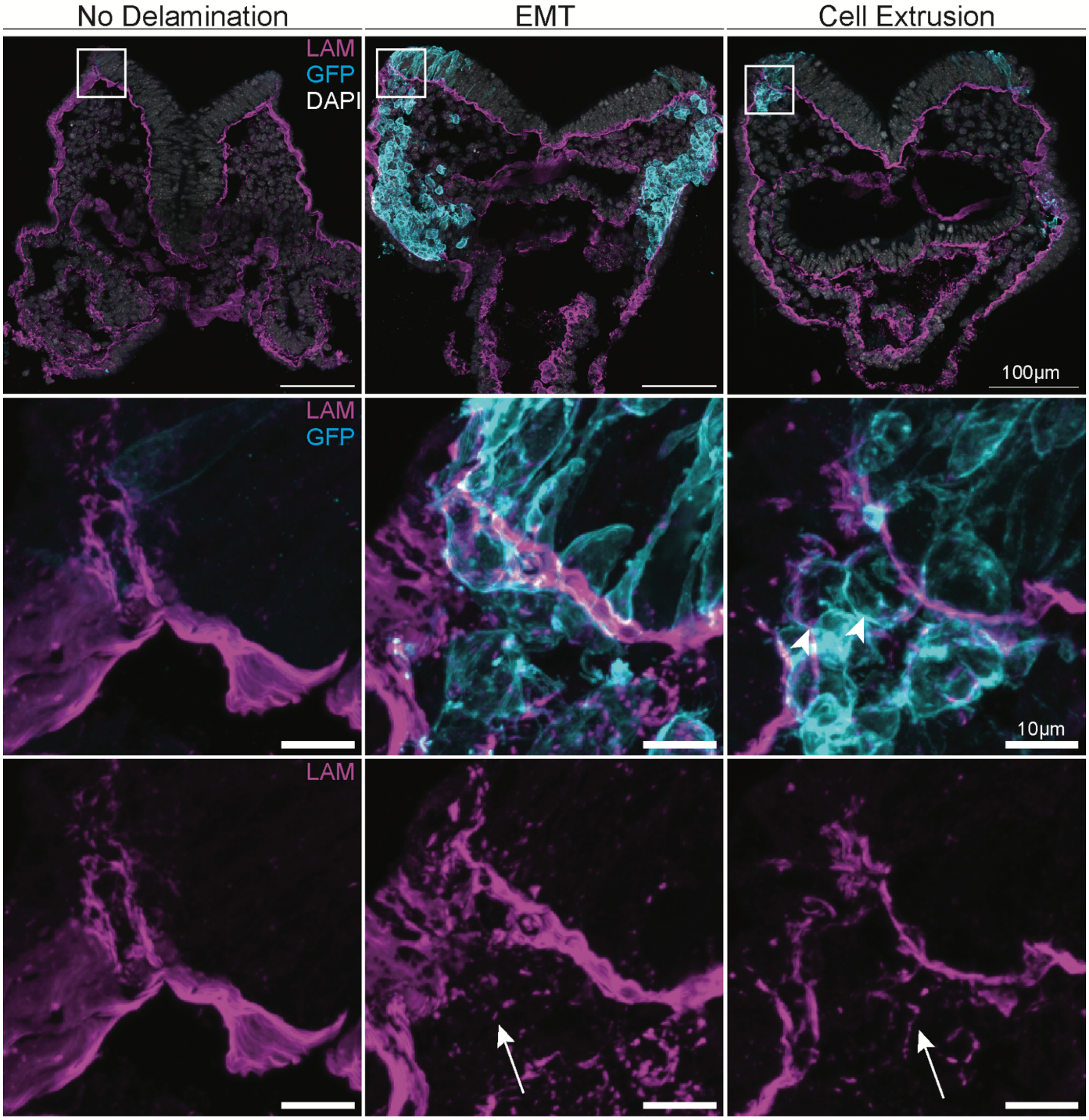


**Supplemental Figure 2. Laminin breakdown has begun by delamination of the round subpopulation.** Immunostaining of Laminin and GFP along with DAPI on transverse histological sections of *Wnt1-Cre;R26R-mTmG* embryos at E8.5. Each column provides an example of the level of laminin breakdown of stages beginning with regions before delamination has taken place, areas where EMT is underway and locations of the round subpopulation identified as undergoing cell extrusion. Boxes indicate the region of the original tissue section on which the zoomed-in images are focused. Arrows point to fragmented laminin in the mesoderm. In regions of tissue lacking NCC delamination, laminin is present in thick structures along the basal side of the epithelium. Tissue sections that exhibited EMT-like delamination of NCC had fragmented laminin throughout the mesenchyme where NCC migration had taken place. In the round delaminating population, some laminin fragments could be found similar to EMT-like delamination, although in observably decreased quantities.

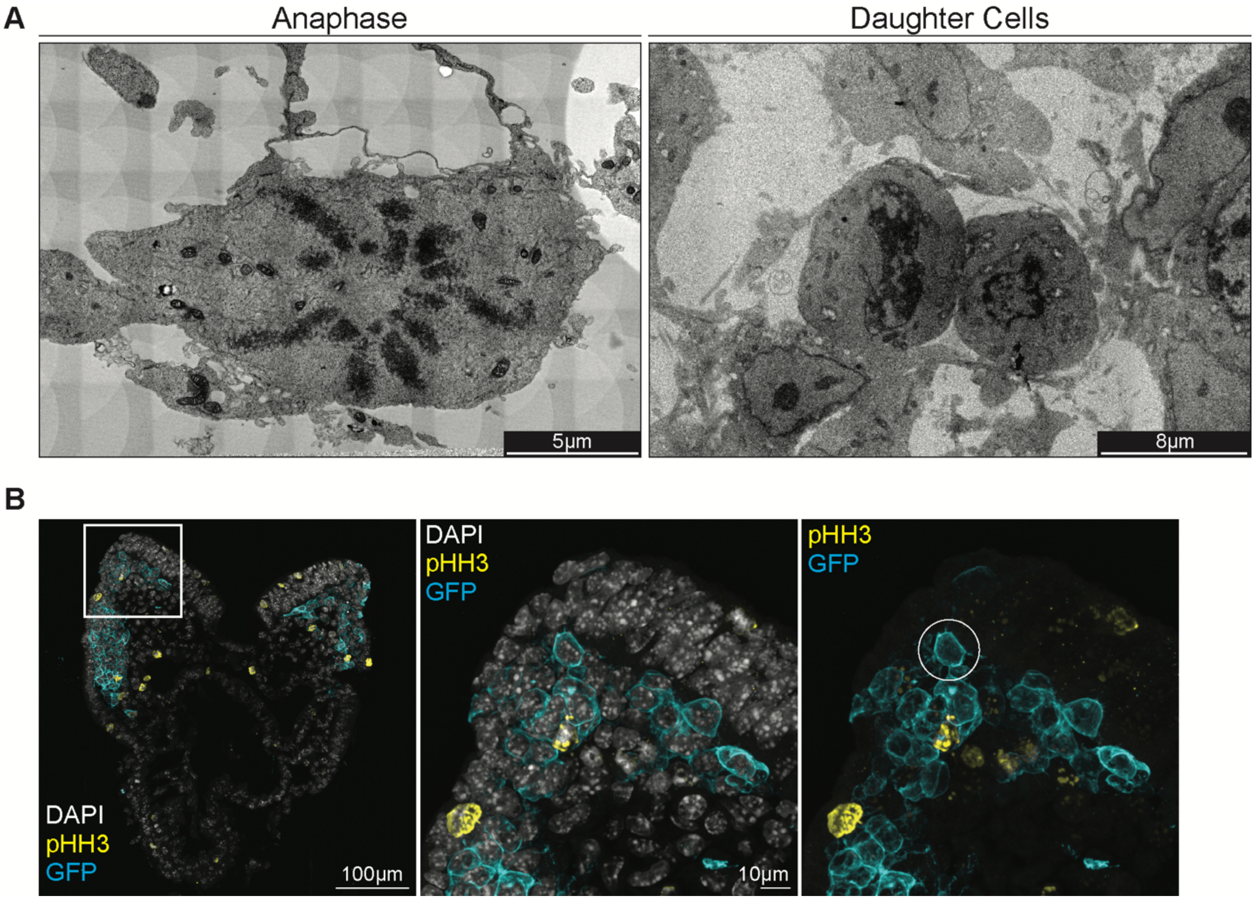


**Supplemental Figure 3. The subpopulation does not appear to be caused by cell division. A**. Transmission electron microscopy on sagittal sections of CD1 (wildtype) embryos at E8.5. Images show the typical chromatin morphologies of cells at various stages of cell division. **B**. Immunostaining of GFP and pHH3 along with DAPI on transverse histological sections of *Wnt1-Cre;R26R-mTmG* embryos at E8.5. The box indicates where the zoomed in images are found in the original tissue section. The circle labels the round NCC which does not express pHH3.

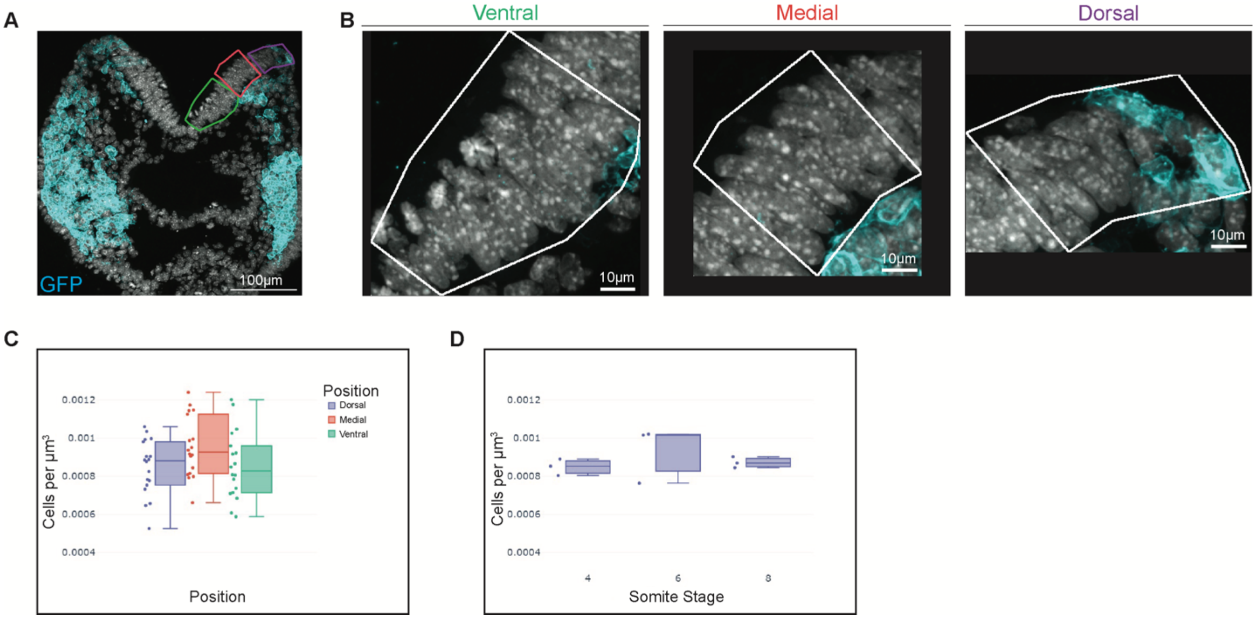


**Supplemental Figure 4. Cell density levels are maintained across the neuroepithelium during NCC development. A**. A max projection of a transverse histological section from a *Wnt1-Cre;R26R*-mTmG embryo at E8.5 used for cell density measurements across the neuroepithelium. The purple region of interest is dorsal, red is medial and green is ventral. **B**. Zoomed in images of the regions of interest from which the cell densities were calculated. **C**. The calculation of cellular density at each region of interest from 3 embryos stages 4-8 somites. No statistical difference was found between all three regions indicating there are no changes in cell density across the neuroepithelium. **D**. The calculation of cellular density across the neuroepithelium from 3 sections each of embryos stages 4-8 somites. No statistical difference was found between all three stages indicating there are no changes in cell density across stages of development.

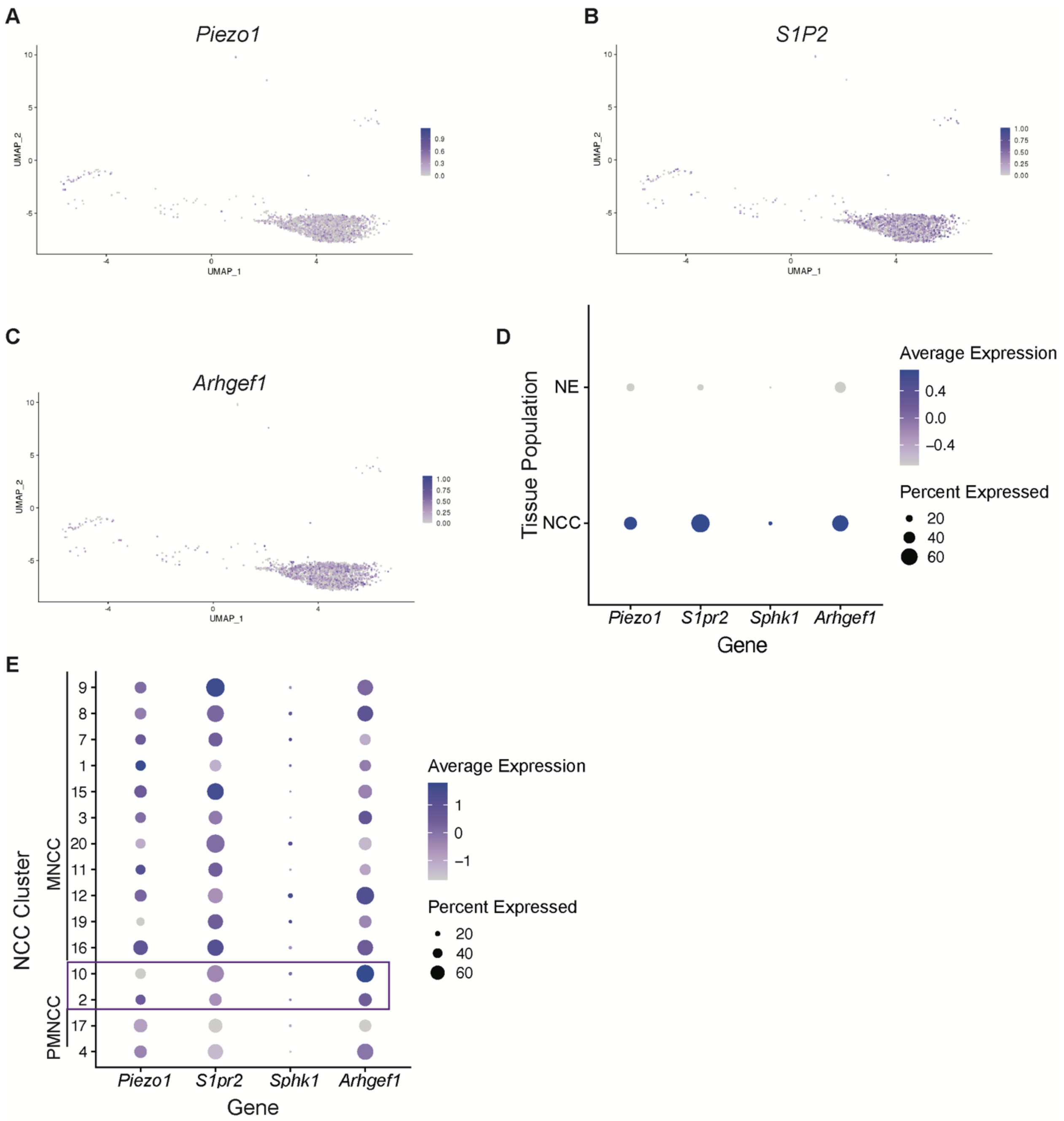


**Supplemental Figure 5. Single cell RNA-sequencing determines live cell extrusion pathway components are expressed during NCC development. A-E**. Expression based on single cell RNA-sequencing of the cranial region of *Wnt1- Cre;R26R-eYFP* and *Mef2c-F10n-LacZ* embryos at E8.5. **A-C**. Expression of *Piezo1*, *S1P2* and *Arhgef1* in the NCC population cluster indicate these cell extrusion factors are expressed in NCC during development. **D**. Dot plot comparison of the NCC compared to neuroepithelium (NE) cells suggests the cell extrusion regulators are specifically expressed in the NCC as compared to neuroepithelial cells. **E**. Dot plot of expression across each stage of NCC development. PMNCC denotes the pre-migratory NCC clusters, the box highlights the delaminating NCC and MNCC labels the migratory NCC populations. These regulators are expressed in the delaminating (box) NCC population. Notably, Piezo1 is specifically enriched in population 2 of the transition NCC.

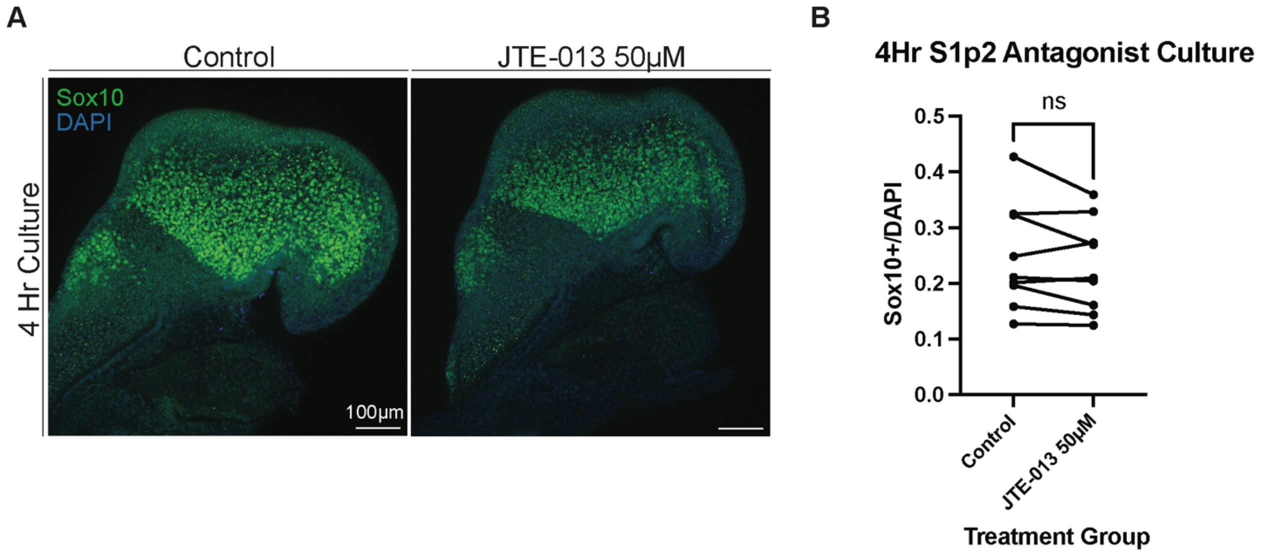


**Supplemental Figure 6. Disruption of NCC delamination is specific to basal live cell extrusion signaling. A**. Immunostaining of Sox10 and DAPI on CD1 embryos at E8.5 following 4-hour culture with drug vehicle (control) or 50µM JTE-013. **B**. Stage matched comparison between control and 50µM JTE-013 embryos of the total number of migratory NCC (Sox10+) normalized to the total number of cells in the head (DAPI). A t-test determined there was no significant difference between S1P2 inhibitor treated and control embryos suggesting S1P2 does not function in NCC extrusion delamination.
